# Deep Learning Genome-wide Linkage Association Study for Wheat Fusarium Head Blight Resistance Genes Discovery

**DOI:** 10.1101/2021.10.11.463729

**Authors:** Wayne Xu, Andriy Bilichak, Raman Dhariwal, Maria A. Henriquez, Harpinder Randhawa

## Abstract

**Background:** *Fusarium* head blight (FHB) is one of the most devastating diseases of wheat worldwide and artificial intelligence can assist with understanding resistance to the disease. Considering different sample populations, marker types, reference maps, and statistical methods, we developed a Deep Learning Genome-wide Linkage Association Study (dpGLAS) of FHB resistance in wheat.

**Results:** The dpGLAS was first applied to two bi-parental population datasets in which the cultivar AC Barrie was a common parent for FHB resistance. Eight candidate gene markers were discovered in the one AC Barrie population and 10 in the other associated with FHB resistance. Eight of these markers were also supported by the conventional QTL mapping. Most of these candidate marker genes were found associated with the Reactive Oxygen Species (ROS) and Abscisic acid (ABA) axes. These ROS and ABA pathways were further supported by RNA-seq transcriptome data of FHB resistant cv. AAC Tenacious, a parent of the third bi-parental population. In this dataset, the ROS-centered Panther protein families were significantly enriched in those genes that had most different response to FHB when compared the resistance Tenacious and the susceptible Roblin.

**Conclusions:** This study developed the framework of dpGLAS and identified candidate genes for FHB resistance in the Canadian spring wheat cultivars AC Barrie and AAC Tenacious.

## Introduction

Modern crop breeding programs are increasingly using genomic selection (GS) and expected to employ gene editing (GE) in the future. Finding the genomic variations that are associated with desirable phenotypes is a crucial step for either GS or GE. Genome-wide association studies (GWAS) and quantitative trait locus (QTL) mapping have been employed to study the genetic basis of numerous traits in the past. With respect to the enormous data generated from different sample populations, types of variations, reference maps, and statistical methodologies, a different analytical design is required to gain new useful information.

Natural population samples of random-mating are commonly used for GWAS[1].GWAS is unable to detect associations between rare allele variants and phenotype because these rare variants appear in a very low frequency in the population[2-4]; however, rare variants may have large effect on the trait of interest[4]. Surprisingly, the rare variants have been found to be more abundant than common variants in the human genome[5]. Bi-parental crosses are commonly used for QTL mappings[6], but the precision for mapping QTLs is typically low and a significant QTL may span a large chromosome region which limits the identification of candidate genes. For example, the confidence interval of a QTL may span 10-20 cM (tens of Mb) and include a chromosome segment containing hundreds of genes[7]. For the large polyploid plant genomes such as wheat genome, most regions of the genome are non-coding regions which are dispersed between coding genes and more than 85% of these non-coding regions consist of repetitive sequences[8]. These non-coding interval regions from conventional QTL mappings are not our first focus. On the other hand, the bi-parental populations are not appropriate for conventional GWAS because they do not comply with the GWAS assumption that the recombinant events are random in the natural population. Also, if the SNP markers based on reference alignment were used in the individual or combined bi-parental population data, the information of the parent genotype was lost who carried the trait of interest. Additionally, the site with different genotypes that both differ from the reference will be labelled as the same marker values.

Types of genetic markers used and the reference genome availability also affect the choice of gene discovery methods. RFLP (restriction fragment length polymorphism)[9], AFLP (amplified fragment length polymorphism)[10], SSRs (microsatellites or single sequence repeats[11], and DArT (diversity arrays technology)[12] markers were developed and used for creating linkage maps and QTL linkage mapping studies particularly when a physical genome map was not available. These non-genic but genomic markers were used intensively in early times before new sequencing technologies emerged. Recent advances in next generation sequencing (NGS) technology enable us to detect genome-wide genetic variants, such as single nucleotide polymorphisms (SNPs) and insertion/deletions (INDELs) at a low cost[13]. The SNPs discovered by mapping RNA-seq from a large number of wheat lines on wheat reference transcripts were used as Microarray beadChip SNP probes and this beadChip platform can detect almost genome-wide genic markers[14]. The whole genome sequencing (WGS) and genotyping-by-sequencing (GBS) which included both genic and genomic SNP markers aid the fine mapping of GWAS and QTLs[15]. The use of a genetic map was the only approach to identify marker locus in QTL linkage mapping before the physical reference genome became available; however, the genetic map is still being used even if the relevant reference genome is available based on the types of sample population in the experiment.

There are a variety of statistical methods implemented in GWAS and QTL linkage mappings. When testing a single locus in a case-control phenotype GWAS, a significance test such as Chi-squared test, odds ratio test, or Fisher’s Exact test is applied to the variant frequency between the two case groups[16]. If the phenotype is a continuous variable, ANOVA[17], t-test[18], or linear regression[19] can be used to evaluate the relationship between the variant types and the phenotype values. When potential confounding variables need to be controlled such as age, gender and medication, the generalized linear model (GLM) can be applied[20]. GWAS has limitation to detect the site interaction or epistasis effect[4]. QTL linkage mapping was designed for bi-parental or multiparent population data[21]. A linear regression model can be applied to single genotype marker association analysis[22]. This genotype association in QTL mapping differs from the variant association analysis in GWAS, even though the same regression model can be used. The single locus mapping results in large genomic regions. Therefore, there is a motivation to scan smaller intervals between markers. This interval mapping approach is to examine how the likelihood ratio changes at different positions along the chromosome, for example, every 2 cM, as a graphical profile[23]. The LOD score (the logarithm of the odds ratio) is a measure of the strength of evidence for the presence of a QTL at a particular location[24]. The epistatic effects of QTLs can be detected by multiple regression models with terms for pairwise interactions between markers[25].

Though the above statistical methods have been well established for GWAS and QTL mapping, application of deep learning could obtain inconceivable outcomes by learning data features[26]. Recently, deep learning has been applied in genetic association studies[27-29]. This artificial intelligent (AI) approach opens a new avenue for genetic association studies. Typically, a neural network uses continuous or discrete values as inputs to predict either a regression or classification output. Ma et al (2018)[27] developed a convolutional neural network (CNN), called DeepGS, to predict phenotypes using SNP markers. A natural population of 2000 Iranian bread wheat landrace accessions were genotyped with 33,709 DArT markers. For the DArT markers, an allele was encoded by either 1 or 0, to indicate its presence or absence, respectively. Liu Y et al (2019)[29] proposed two-stream CNN model for bi-parental population and genomic SNP data. But it did not focus on the parent genotype of the trait of interest and it does not work for dataset integration. Further, it did not thoroughly search for the spatial effect via different combinations of gene marker orders.

FHB is one of the most devastating disease of wheat worldwide, leading to severe yield and quality losses[30]. The genetic basis of FHB resistance in wheat has been studied extensively, and more than 100 QTL associated with FHB resistance have been reported on the 21 wheat chromosomes[31]. Among these FHB-resistant QTLs, *Fhb1*[32], *Fhb2*[33] and *Fhb5*[34] on chromosome arms 3BS, 6BS, and 5AS, are the most-characterized. *Fhb1* (*QFhs*.*ndsu-3BS*) on chromosome arm 3BS, derived from cultivar Sumai-3 and its derivative Ning 7840 was described as the strongest and best-validated FHB-resistance QTL and is primarily associated with type II resistance[35, 36]. However, these loci spanned a large region in cM scales. It is challenging to fine map and identify the functional genes in these loci, though recently a pore-forming toxin-like (PFT) gene[37] and a putative histidine-rich calcium-binding protein (TaHRC)[38] were reported as the genes responsible for FHB resistance at the *Fhb1* locus. Despite these extensive works, little is understood about the genetic basis of native FHB resistance in Canadian spring wheat (i.e., FHB resistance not introduced from Sumai-3 and other Asian spring wheats). A recent study of the AC Barrie, a hard red spring wheat cultivar in the Canada Western Red Spring marketing class that possesses an intermediate level of FHB resistance, by QTL linkage mapping discovered 21 QTLs associated with FHB resistance[39]. However, it is far away for gene identification.

We propose here a Deep Learning Genome-wide Linkage Association Study (dpGLAS) strategy for FHB resistance gene discovery. dpGLAS takes into consideration the types of sample populations, marker features, reference map availability, and the analytic methodologies. dpGLAS is different from GWAS in that it uses genotype information of the parental population to determine marker candidates instead of using the variants, such as SNPs. Moreover, it differs from conventional QTL mappings in that dpGLAS uses genome reference and all co-segregating representatives and is not interested in searching those non-coding loci between functional sites. Whereas conventional QTL mapping uses genetic linkage map and marker binning. Neural network deep learning replaced other statistics mining methods in the QTL mapping and GWAS. We applied dpGLAS to understand the FHB resistance in a Canadian spring wheat cultivar, AC Barrie, which is a hard red spring wheat cultivar in the Canada Western Red Spring marketing class that possesses an intermediate level of FHB resistance.

## Results

### Genome-wide genic markers covered

The probes in the Illumina Infinium 90K wheat SNP beadchip were previously derived from the RNA-seq transcriptome of wheat which includes about 90,000 gene-associated SNPs[14]. This number is close to the wheat gene number of 108,639 covered by a comprehensive RNA-seq transcriptome data[40]. As shown in **Table 1**, approximately 75% of the parental genotype marker sites were identical between the FHB resistant parent (B) and susceptible parent (A) genotypes which reflected monomorphic sites in these bi-parental populations. These monomorphic markers were removed from analysis because these sites are not of interest. Marker sites ranging from 4917 to 11336 had identical variants (co-segregating markers) in populations. The remaining ranging from 1892 to 5872 genotype markers further retained by samples with trait information were worth investigation. The 1892 to 5872 detectable markers had 4917 to 11336 co-segregating markers (**Table 1**). As such, the detectable candidates can be used to trace or fine map to a scale of 1-5 sites on average. About 6,000-10,000 (∼10%) marker sites were removed due to missing in parent genotyping or population genotyping. As a result, more than 70,000 sites were covered by the analytic design which represented most of the genes of interest in the whole transcriptome.

**Table 1.**
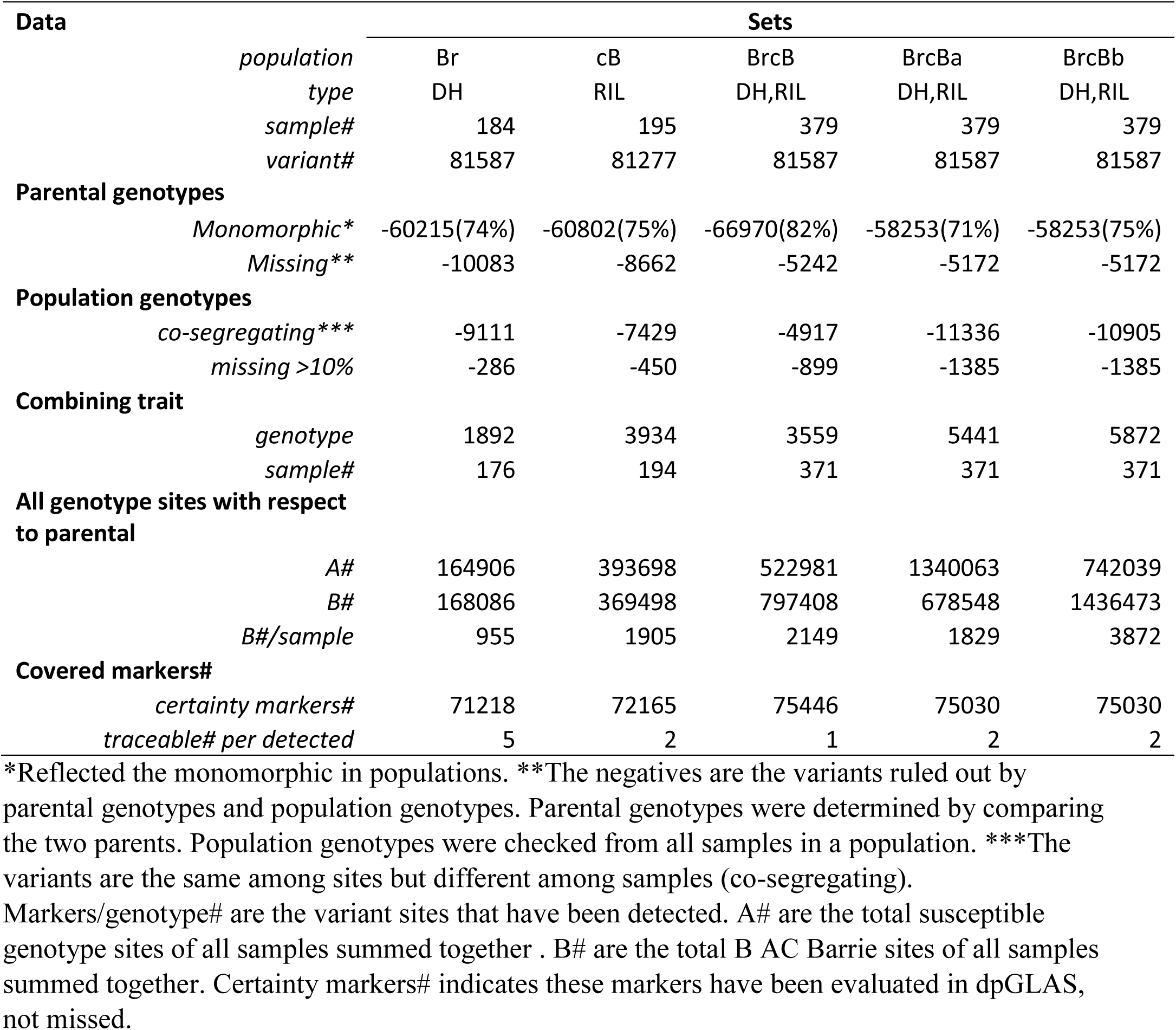
Gene markers and filtering in five datasets

| Data | Sets |  |  |  |  |  |
| --- | --- | --- | --- | --- | --- | --- |
|  | *population* | Br | cB | BrcB | BrcBa | BrcBb |
|  | *type* | DH | RIL | DH,RIL | DH,RIL | DH,RIL |
|  | *sample#* | 184 | 195 | 379 | 379 | 379 |
|  | *variant#* | 81587 | 81277 | 81587 | 81587 | 81587 |
| **Parental genotypes** |  |  |  |  |  |  |
|  | *Monomorphic\** | -60215(74%) | -60802(75%) | -66970(82%) | -58253(71%) | -58253(75%) |
|  | *Missing\*\** | -10083 | -8662 | -5242 | -5172 | -5172 |
| **Population genotypes** |  |  |  |  |  |  |
|  | *co-segregating\*\*\** | -9111 | -7429 | -4917 | -11336 | -10905 |
|  | *missing >10%* | -286 | -450 | -899 | -1385 | -1385 |
| **Combining trait** |  |  |  |  |  |  |
|  | *genotype* | 1892 | 3934 | 3559 | 5441 | 5872 |
|  | *sample#* | 176 | 194 | 371 | 371 | 371 |
| **All genotype sites with respect to parental** |  |  |  |  |  |  |
|  | *A#* | 164906 | 393698 | 522981 | 1340063 | 742039 |
|  | *B#* | 168086 | 369498 | 797408 | 678548 | 1436473 |
|  | *B#/sample* | 955 | 1905 | 2149 | 1829 | 3872 |
| **Covered markers#** |  |  |  |  |  |  |
|  | *certainty markers#* | 71218 | 72165 | 75446 | 75030 | 75030 |
|  | *traceable# per detected* | 5 | 2 | 1 | 2 | 2 |

\*Reflected the monomorphic in populations. \*\*The negatives are the variants ruled out by parental genotypes and population genotypes. Parental genotypes were determined by comparing the two parents. Population genotypes were checked from all samples in a population. \*\*\*The variants are the same among sites but different among samples (co-segregating). Markers/genotype# are the variant sites that have been detected. A# are the total susceptible genotype sites of all samples summed together. B# are the total B AC Barrie sites of all samples summed together. Certainty markers# indicates these markers have been evaluated in dpGLAS, not missed.

The integration of the two datasets increased the population size to 371 samples. The detectable genotype marker numbers of the integrated datasets, BrcBa (5,441) and BrcBb (5,872), were higher than the independent sets, Br (1,892) and cB (3,934), though BrcB had 3,559. The total marker loci carrying the B genotype and loci per sample were different among the single sets and integrated set (**Table 1**).

### Optimized NN models and validations

The Br dataset was used for the optimization of deep learning NN architecture and parameters. The input features were genotype marker binary values and the predict traits were the continuous values of FHB incidence and severity rating index (VRI). The training models using a 0.9 portion of the samples applied to the 0.1 portion of samples as a test set predicted the original FHB trait with a mean correlation of 0.42 (std 0.14) (Supplemental **Table S1)**. The one convolution (Conv) layer plus a Dense layer (CNN11) had a lowest performance (mean 0.352, std 0.177). When increasing the Conv stream (CNN22), the prediction correlation increased. The CNN22 was selected for conducting further parameter optimization. Six parameters were evaluated for three repeat tests. “TruncatedNormal” and “RMSprop” were chosen to be the initializer and optimizer, respectively. “Linear” activation was chosen for both Conv and Dense layers. The filter size had a combination of 6,10,4 with the filter pool size of 2 (Supplemental **Table S2)**.

The optimized NN models were applied to five datasets. After 10 repeat updates (epochs), the loss of mean squared error (MSE) dropped to 0.2 in the DNN learning (**Fig 2a**). After 60 epochs, the MSE approached zero and stabilized. The SHAP scores obtained at this stable stage allowed the ranking of importance of gene markers. In the same dataset, the markers with relatively high scores had high frequency in the top list, though they may not have the exact same order. The top 100 markers were assumed to be enough or more than the associated markers for FHB resistance and these markers had best explained the trait of the whole data sample set.

**Fig 1.**
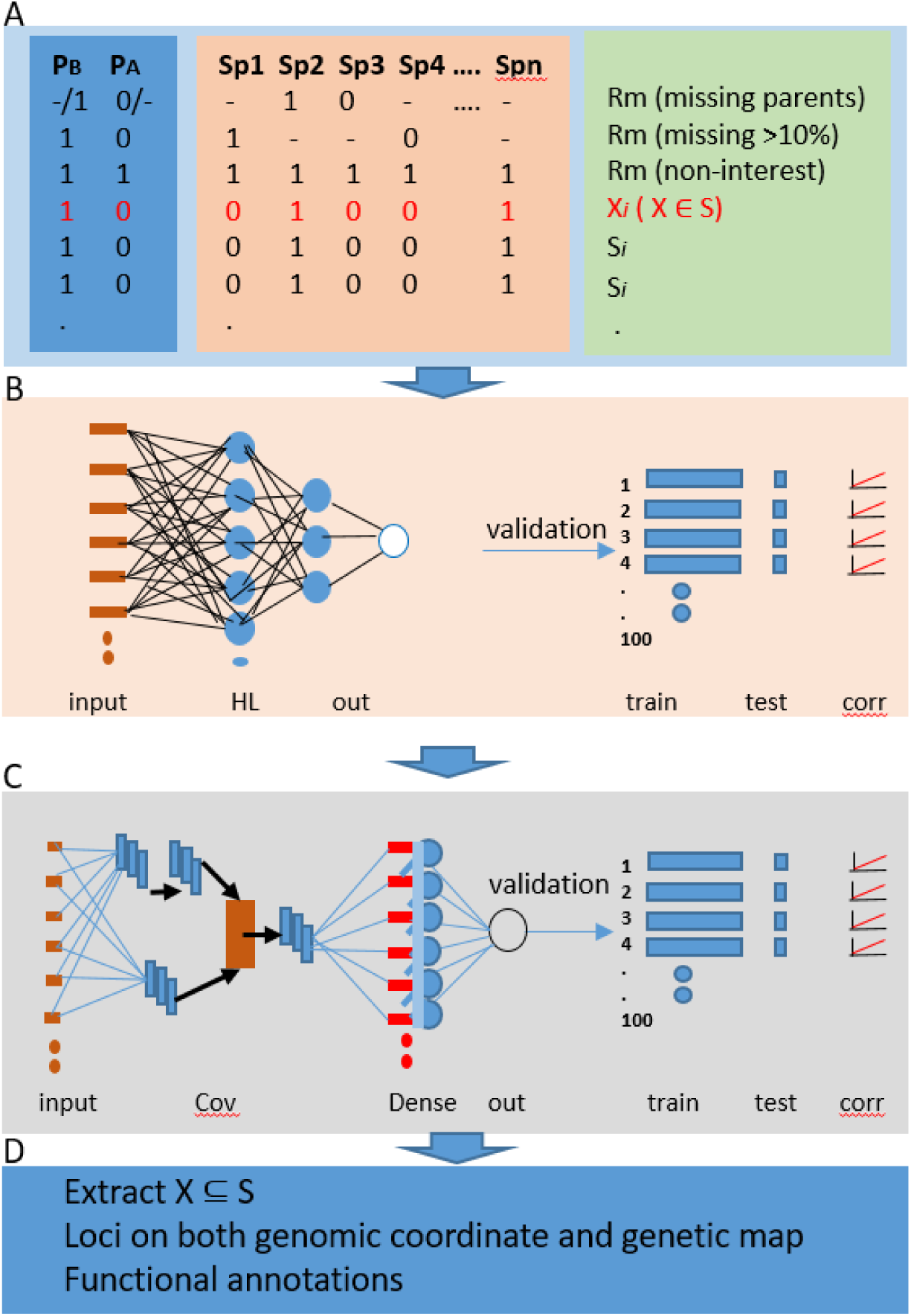
dpGLAS schema. A. marker genotype, linkage, and integration (P_B_: genotype of parent AC Barrie. P_A_: genotype carried by the other parent either Cutler or Reeder. Sp1, Sp2,.. :sample 1,2,.. in each bi-parental populations. X_*i*_: site *i* belongs (ϵ) to one of the co-segregating set, S_*i*_: set *i* of co-segregating). B. DNN step with 10-fold validations (HL: hidden layers). C. CNN step with 10-fold validations (Conv: Convolutional layers; Dense: Dense layers). D. candidate gene annotation.

**Fig 2.**
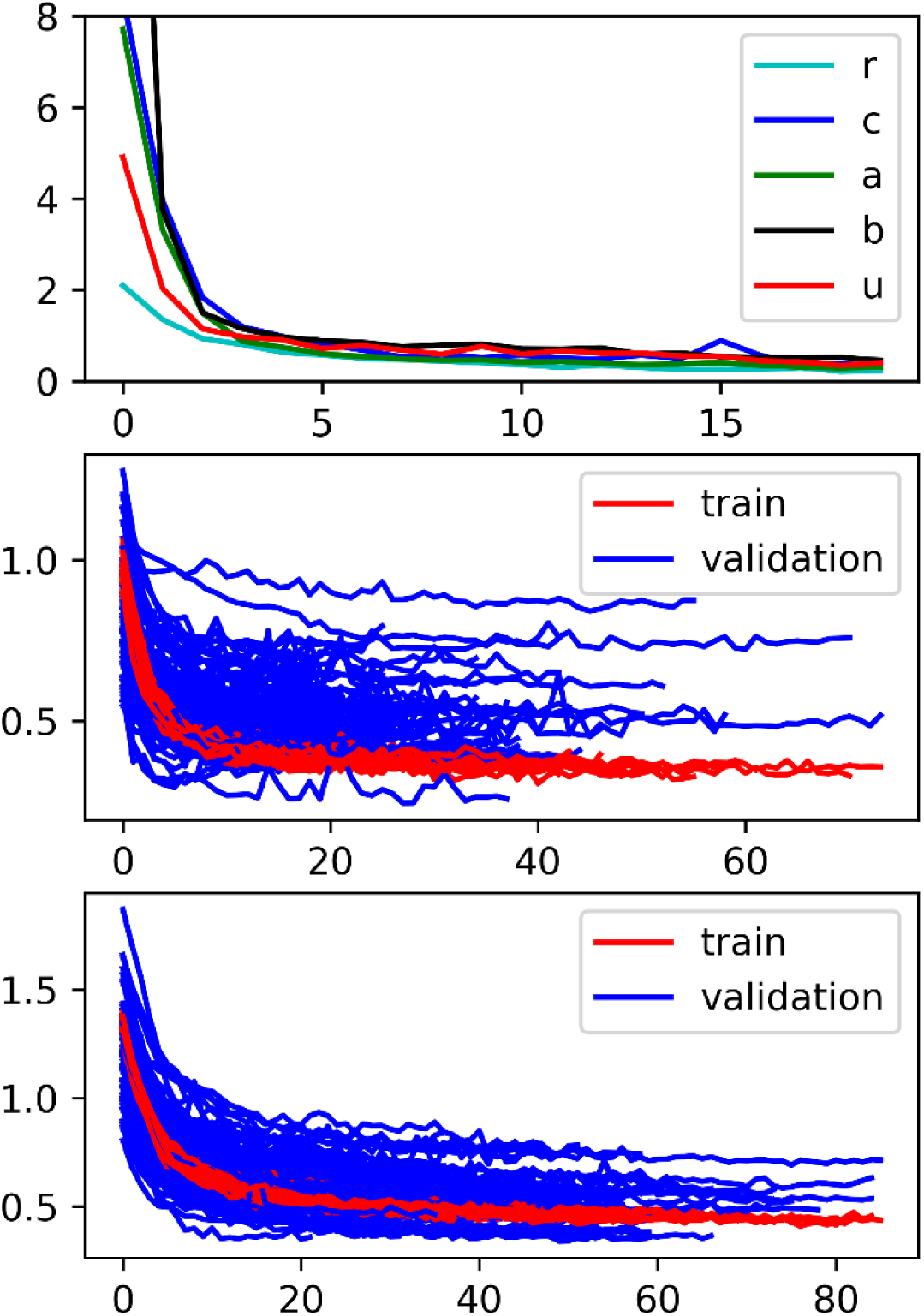
Loss curves of mean squared errors (MSE) of models. A. MSE loss (approaching) to zero using the whole dataset of the five sets. B. MSE loss of training samples (0.9, red) and test samples (0.1, blue) in 100 DNN model runs of the BrcBa integrated set. C. MSE loss of training samples (0.9, red) and test samples (0.1, blue) in 100 CNN model runs of the BrcBa integrated set. One letter represents dataset: r: Br; c:cB; u: BrcB; a: BrcBa; b:BrcBb.

In the 100 times of 10-fold validations, the weights of the 0.9 portion of samples (training set) were re-evaluated to generate a new model and this newly-generated model was applied to the 0.1 portion of test set (validation set). The MSE curves demonstrated the model performance and that the MSE curves of test set (blue) had a similar pattern and trend to those of the training set (red) (**Fig 2b**, Supplemental **Fig S1**), with only few over-fitting (curves under the red curves) and under-fitting (curves above the red curves). For example, in BrcB set, the final MSE scores of training sets and tests were 0.117 (std 0.0123) and 0.2781 (std 0.096), respectively. In these 100 times of validations, the SHAP scores and prediction correlations were examined. A random 10-fold validation test of BrcB by DNN model showed a 0.61 Pearson correlation score (**Fig 3a**). The average prediction correlation score of 100 random tests was 0.70 (std 0.07) for BrcB (u in red, **Fig 3c**).

**Fig 3.**
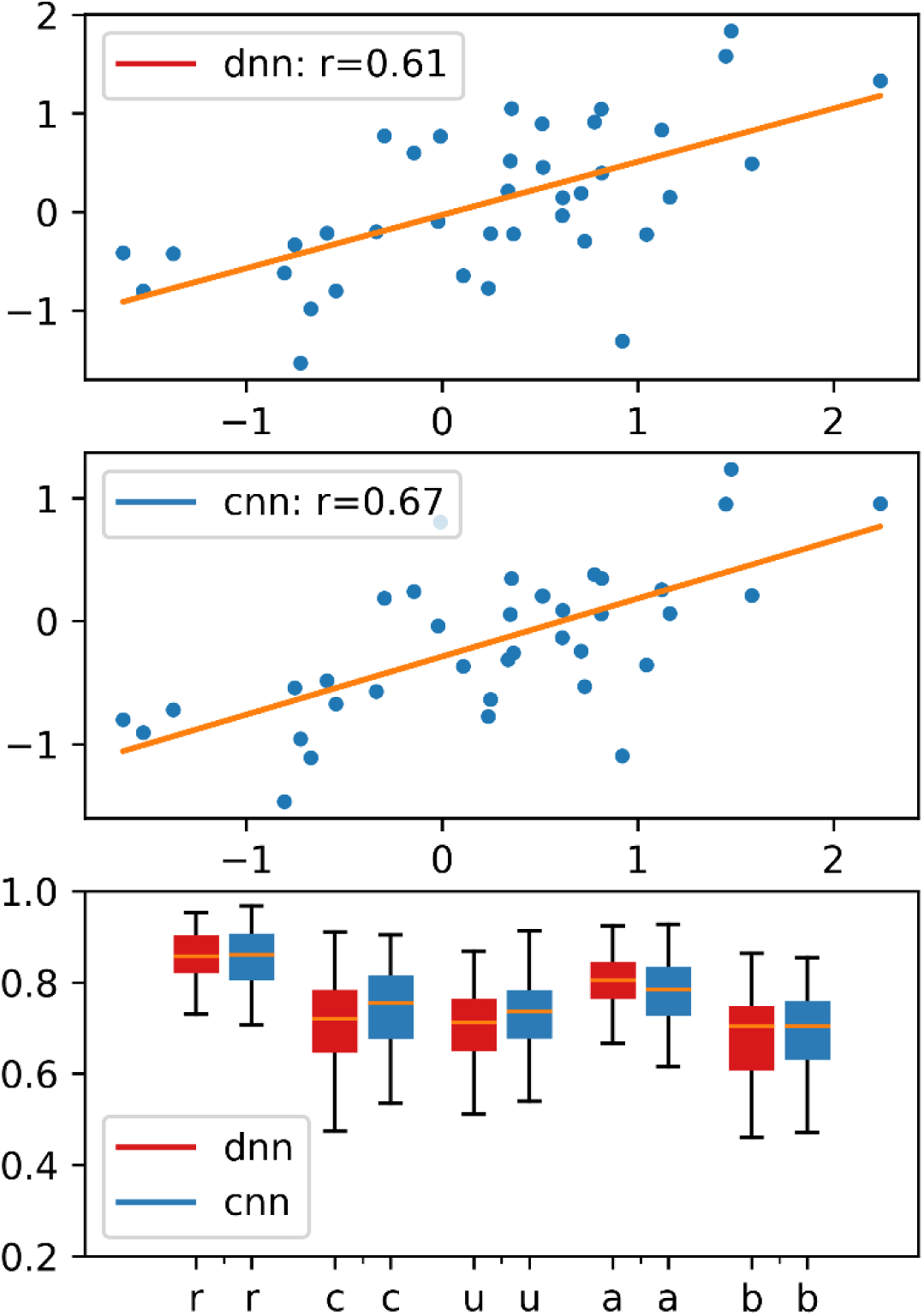
Pearson correlation between the original FHB infection visual rating index (VRI) and the predicted VRI values. A. a random test of DNN model using BrcB set. B. the same random samples but tested by CNN model. C. the overall correlations in 100 runs of all five datasets (One letter represents datasets: r: Br; c:cB; u: BrcB; a: BrcBa; b:BrcBb).

The spatial epistatic effects among the 100 gene markers were detected in the whole sample set using CNN model. The CNN ran 100 times and each time the marker order was updated by the SHAP scores. The marker order was saved as the best one when the sum of the SHAP scores of the top 20 markers reached the highest. The top 20 markers from three sets are shown in **Fig 4**. This saved marker spatial order was examined 100 times using 10-fold validation. As shown in **Fig 2c**, the CNN model followed DNN step had a better performance over DNN model alone in **Fig 2b** in that the test MSE curves were more converged to training curves. A 10-fold validation test of the CNN model for the same BrcB test set had a higher Pearson correlation score (0.67) than DNN alone (**Fig 3b**). The average prediction correlation score of 100 random tests was 0.73 (std 0.08) for BrcB (u in blue, **Fig 3c**).

**Fig 4.**
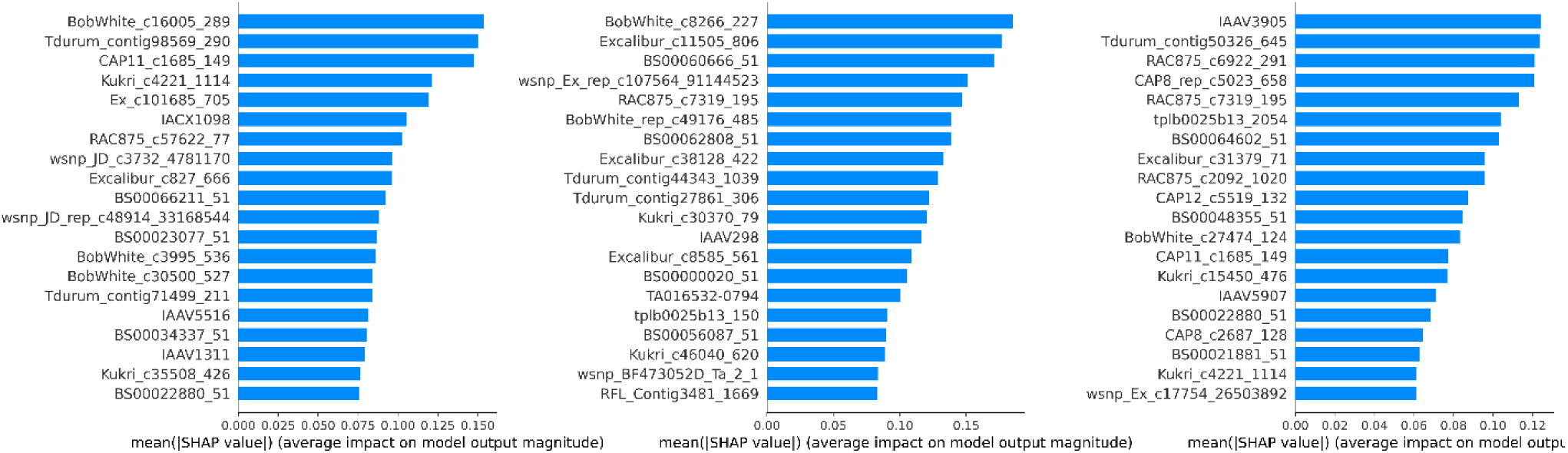
Gene marker ranking by SHAP importance scores in dataset Br (A), cB (B), and the integrated BrcBa (C). The SHAP Scores were generated by CNN model from the whole dataset. The marker order was saved as the best one when the sum of the top 20 markers’ SHAP scores reached the highest. The marker orders will be examined 100 times using 90% subsets to determine the markers that appear at least 80 times in the top 10.

### Top associated markers and Integrative analysis

In Br set, eight variants/groups (**Table 2)** appeared in the top 10 list with >80 times in the 100 validations of either DNN (Pearson correlation rate of 0.85, **Fig 3c**) and/or CNN (Pearson correlation rate of 0.85). In cB set, 10 variants/groups were found that appeared in the top 10 list with >80 times in the 100 validations of either DNN (Pearson correlation rate of 0.69, **Fig 3c**) and/or CNN (Pearson correlation rate of 0.72). The two different types of parental populations (DH and RIL) had the same parent genotype B (AC Barrie). But there was no overlap between these two lists. Only one overlap was found from these two datasets when we selected the top 20 markers that best fitted the DNN models in each dataset alone (Supplemental **Fig S2a)**, and the overlap increased to 3 markers when the spatial effects were considered (Supplemental **Fig S2b)**. To look into the cause of the low overlap rate between these two sets, we found that the whole 11,003 (9,111+1,892, **Table 1**) markers of Br and 11,363 (7,429 + 3,934) of cB only had around one third (4,147) in common because of monomorphic and missing in either one of these two sets. In the final discovered lists (**Table 2**), 5 variants/groups were missed in another set because of monomorphic, therefore a high rate of common variant sites were not seen from the Br and cB sets even though they had the common AC Barrie parent.

**Table 2.**
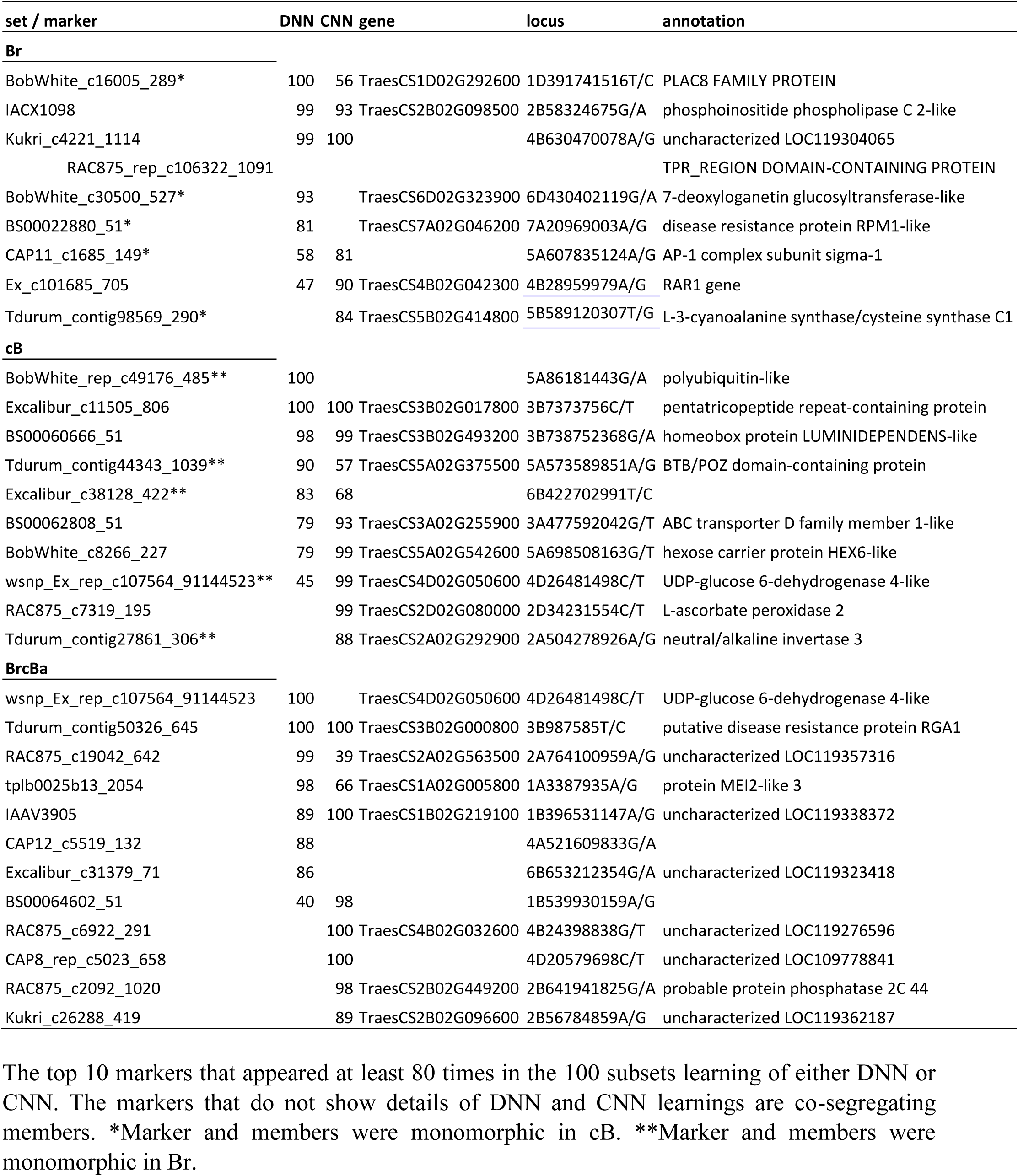
The top marker associated genes and annotations from three set analyses.

Combining sets into larger sample size had a few more overlaps compared to each individual set (BrcB, Supplemental **Fig S2 c&d**), indicating that BrcB allowed the detection of those markers that were not in the independent set test alone. The overlaps of the three different integrations were shown (Supplemental **Fig S2ef)**.

The loss of MSE curves showed that all five datasets had a similar model performance in terms of training and test curves, but BrcBa had a better performance in CNN modeling than DNN (**Fig 2**, Supplemental **Fig S1d**). Among the three types of data integrations from the same two independent sets, BrcBa showed the highest prediction correlation score (DNN of 0.79, CNN of 0.78) (**Fig 3c**). BrcBb was the lowest at 0.69 (CNN). These results indicated that the gene markers learned from the combined set were able to explain and predict the traits from different independent sets.

### Candidate genes

In Br set, the top eight variants/groups were located in chr1B, 2B, 4B, 5A, 5B, 6D and 7A (**Table 2)**. Six markers harbored annotated genes while two markers’ loci did show gene and expression in AC Barrie by RNA-seq signals, though no gene annotation (Supplemental **Fig S3**). The Kukri_c4221_1114 marker appeared in the top 10 list of all 99 subset DNN validations and 100 subset validations. This marker and the four co-segregating markers were located in chr5B. It appears that the two bi-parental populations (Br and cB) with the common AC Barrie had a different segregating pattern. The RIL population (cB) had more through segregation with more single member locus than DH (Br) did (Supplemental **Table S3**). One member of Kukri_c4221_1114 group in Br set RAC875_rep_c106322_1091 harbored within gene TraesCS4B02G337300 which encodes a TPR_region domain-containing protein. This variation caused a missense mutation p.Thr130Met in the exon 2 (**Fig 5A**). The marker BS00022880_51 in chr7A:20969003 was also a missense variant (p.Tyr397Cys) and belong to disease resistance protein RPM1.

**Fig 5.**
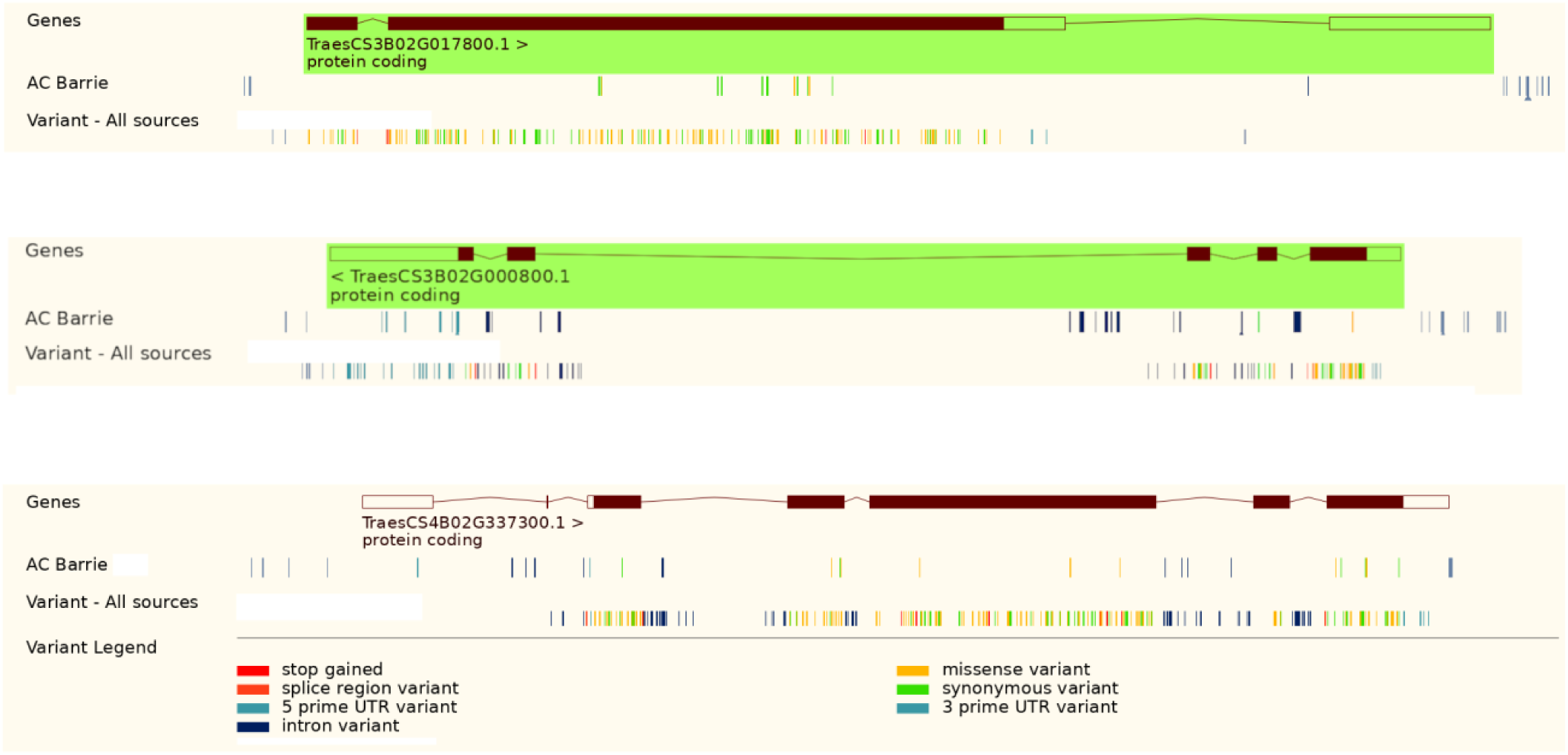
Variations in three genes of AC Barrie and other sources of wheat. AC Barrie variant vcf files were downloaded from AC-Barrie WGS vs IWGSC RefSeq v1.0 Genome Assembly (figshare.com) The IWGSC RefSeq v1.0 was viewed by plant Ensembl browser. “AC Barrie” panel was custom added into browser. “variant – All sources” panel was Ensembl browser built-in.

In set cB, the top10 variants/groups were in chr2A, 2D, 3A, 3B, 4D and 5A (Supplemental **Fig S4**). The topmost one was Excalibur_c11505_806 which repeatedly appeared in 100% validations. Though this was a synonymous variant, p.Ser785Ser, the associated gene TraesCS3B02G017800 encoding pentatricopeptide repeat-containing protein carried many variants compared to other susceptible cultivar Fielder (**Fig 5C**,). RAC875_c7319_195 in chr2D:34231554, at the 3’UTR region of gene TraesCS2D02G080000 encodes L-ascorbate peroxidase 2. The susceptible Fielder did not harbored this variant.

In the integrated dataset BrcBa, 12 variants/groups were found in chr1A,1B, 2A, 2B, 3B, 4A, 4B, 4D, and 6B (Supplemental **Fig S5**). The topmost is Tdurum_contig50326_645 variant which occurred at the 3’UTR of gene TraesCS3B02G000800 in chr3B:987585. This gene encodes putative disease resistance protein RGA1 (cullin-RING ubiquitin ligase). The susceptible Fielder showed the variant at this site.

### A third dataset mapping and candidate gene validation

A QTL mapping DH population dataset with FHB resistant parent AAC Tenacious was tested by dpGLAS and eight variants/groups were found in chr2D, 4A, 5D, 6A and 7B. These variants appeared in the top 10 list repeatedly in more than 80 out of 100 subset validations of DNN and or CNN learning (Supplemental **Table S4**). Ex_c7626_392 variant at gene TraesCS4A02G023700 in chr4A:16967586 encodes myosin-binding protein 3-like and had a missense mutation p.Leu663Ser. However, this gene did not express in the AAC Tenacious RNA-seq sample, though the susceptible Roblin showed a low level expression and the same variant (Supplemental **Fig S6**). Another variant in the same group, Ku_c7594_1179, also had a missense variant, p.Lys490Arg. The gene TraesCS7B02G018300 coding for RGA5-like protein had a variant in intron (Supplemental **Fig S7**). Gene TraesCS2D02G080000 encoding L-ascorbate peroxidase 2 had a variant in intron, which is located within peak position of the most important QTL of AAC Tenacious and very close (< 0.3 mb) to tightly linked marker *Ppd-D1* [41](Supplemental **Table S4**). This gene was also detected in dataset cB. Marker member wsnp_CAP7_c44_26549 on chr7B was overlapped by conventional QTL mapping and RAC875_c7319_195 exhibited epistasis QTL in conventional QTL mapping [41](Supplemental **Table S4**).

Since the main FHB resistance was carried by the cultivar AAC Tenacious, we observed the expression levels of the six candidate genes using the RNA-seq data of AAC Tenacious with FHB inoculation and we calculated the differential response genes (DrGs). The differential responses of the 6 candidate genes to FHB inoculation between AAC Tenacious and Roblin are shown in **Fig 6A**. The expression fold changes after FHB inoculation were similar between AAC Tenacious and Roblin (red versus purple) for three genes, *APX2, MAP6*, and *RGA5*; however, the response fold of AAC Tenacious was higher than Roblin (green versus blue) for *APX2, MAP6*, and reductase. We also searched the topmost genes that had most different response to FHB compared in resistant AAC Tenacious and susceptible Roblin. A total of 218 genes were discovered (Supplemental **Table S5)**. Within these genes, the ROS-centered Panther protein families were significantly enriched (**Fig 6B**).

**Fig 6.**
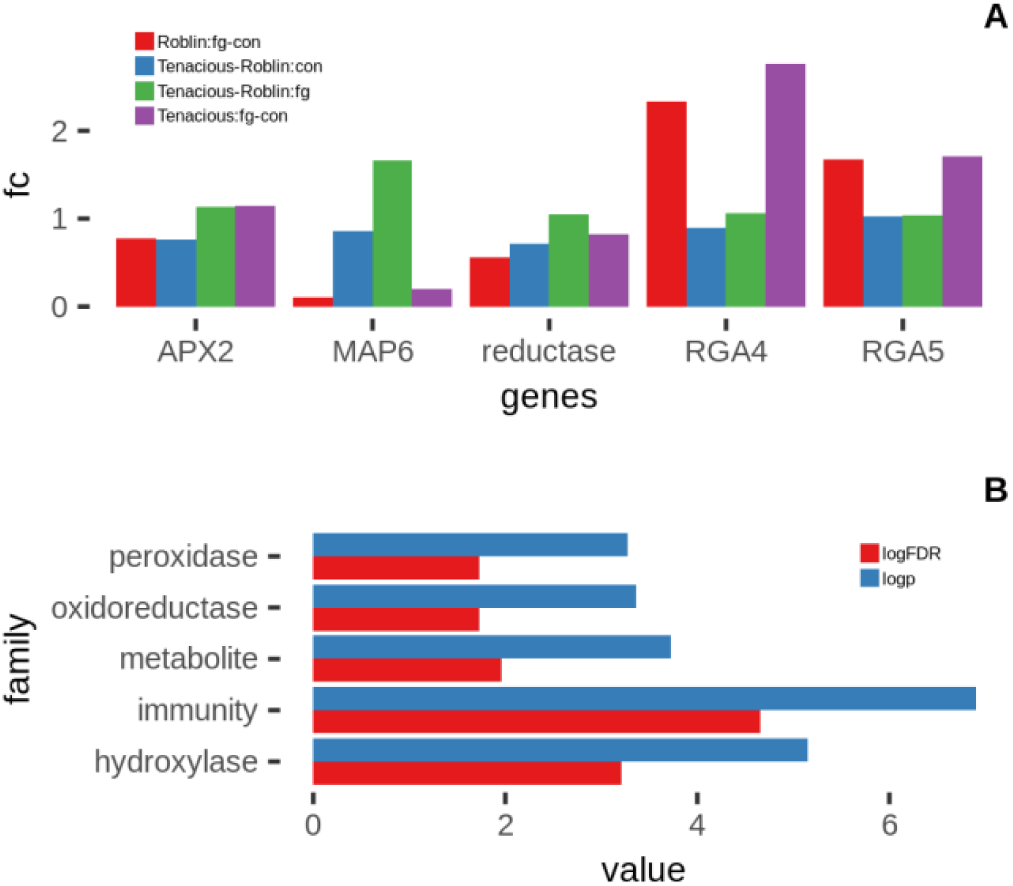
BarChat of RNA-seq of the 6 Tenacious candidate genes. A. the fold change (fc) of two condition comparison: FHB inoculation versus control (fg-con), AAC Tenacious versus Roblin (Tenacious-Roblin). B. Panther protein family enrichment analysis using 215 genes that were the top differential response to FHB (DrGs).

### dpGLAS compared to conventional QTL mapping

As shown in **Table 3**, the QTL linkage mapping resulted in 7 candidate QTL loci from Br population which spanned 1-9 cM range and 4 QTL loci from cB population spanned 1-5 cM at the LOD cutoff of 2.5. In the 100 repeats of QTL mapping using subsets, some QTLs were not detected in subsets. The locus with LOD of 28.02 was missed 13 times out of 100. However, there were 5 marker overlaps between QTL mapping and dpGLAS in Br set and 3 overlaps in cB set when the LOD 2.5 was set for QTL mapping and top 10 was set for dpGLAS. With multiple bi-parental datasets available and large number of genic markers covered, dpGLAS had advantages in the data integration, validation, fine mapping, and gene annotation. These gene annotations allow for further functional, interaction, and pathway analysis.

**Table 3.** Overall comparison of dpGLAS and QTL linkage studies.

| QTL mapping |  |  |  |  |  |  |  |  | dpGLAS |  |
| --- | --- | --- | --- | --- | --- | --- | --- | --- | --- | --- |
| chr | Position | LOD | PVE(%) | Add | LeftCI | RightCI | size cM | *subsets | candidate | annotation |
| **Br set** |  |  |  |  |  |  |  |  |  |  |
| 1B | 44.00 | 4.03 | 2.80 | -2.06 | 43.50 | 44.50 | 1.00 | 66 |  |  |
| 2B | 42.00 | 7.28 | 5.27 | -2.86 | 37.50 | 42.50 | 5.00 | 100 | TraesCS2B02G098500 | phosphoinositide phospholipase C 2-like |
| 2D | 85.00 | 4.49 | 3.10 | -2.19 | 84.50 | 86.50 | 2.00 | 96 |  |  |
| 4B | 53.00 | 6.24 | 4.49 | -2.63 | 50.50 | 53.50 | 3.00 | 36 | TraesCS4B02G042300 | RAR1 gene |
| 4B | 81.00 |  |  |  |  |  |  | 10 | TraesCS4B02G337300 | TPR_REGION DOMAIN-CONTAINING PROTEIN |
| 5A | 136.70 | 20.96 | 18.78 | 5.34 | 135.50 | 136.50 | 1.00 | 100 |  | AP-1 complex subunit sigma-1 |
| 5B | 115.00 | 28.02 | 27.31 | -6.43 | 113.50 | 115.50 | 2.00 | 87 | TraesCS5B02G414800 | L-3-cyanoalanine synthase/cysteine synthase C1 |
| 7A | 12.00 | 3.67 | 2.53 | 1.96 | 7.50 | 16.50 | 9.00 | 79 | TraesCS7A02G046200 | RPM1 |
| **cB set** |  |  |  |  |  |  |  |  |  |  |
| 1B | 70.00 | 3.78 | 5.41 | -2 | 69.5 | 74.5 | 5.00 | 22 |  |  |
| 1B | 106.00 |  |  |  |  |  |  | 22 |  |  |
| 2D | 8.00 | 3.67 | 5.05 | -1.9 | 7.5 | 10.5 | 3.00 | 71 | TraesCS2D02G080000 | L-ascorbate peroxidase 2 |
| 3B | 9.00 | 4.07 | 5.74 | -2 | 8.5 | 9.5 | 1.00 | 90 | TraesCS3B02G017800 | pentatricopeptide repeat-containing protein |
| 3B | 90.00 |  |  |  |  |  |  | 14 | TraesCS3B02G493200 | homeobox protein LUMINIDEPENDENS-like |
| 4D | 30.00 | 14.49 | 24.40 | -4.2 | 27.5 | 31.5 | 4.00 | 100 | TraesCS4D02G050600 | UDP-glucose 6-dehydrogenase 4-like |

\*Frequency detected for repeating 100 times of 90% subsets under LOD cutoff of 2.5. The rows that do not show details of information indicated not detected in that method. dpGLAS used top 10 and detected at least 80 times out of 100 in either DNN or CNN learnings.

## Discussion

In this study, we have developed a deep learning framework for Genome-wide Linkage Association Study (dpGLAS). The performance of deep learning evaluated by the Pearson correlation of the predicted to the original values is superior to the correlation score ranges of the production traits of soybean predicted by deepGS and other statistical methods[29]. Deep learning is focusing on the prediction by combining the contributions of all features and these combinations may compromise the rankings of features when repeating the learning experiment. When we were disentangling the learned parameters using a portion (90%) of samples many times, we found that the features that contributed the most to the model appeared in a range of top list, though not appeared in exactly the same order each time. As such, the 10-fold (90% learning samples, 10% test) validation was implemented in the marker/gene discovery. Whereas GWAS or the conventional QTL mapping applies a significance test to the whole population samples for a single p-value or likelihood odd ratio for each marker/gene or interval. The individual p-values were not intervened by all markers as a whole; however, when the sample is changed, for example, when only a subset of samples were used, the p-values in GWAS and QTL mapping will be different. A “Dropout” function and an “EarlyStopping” function were implemented in NN which simplified learning complex thereby overcoming overfitting but may also attribute to the inconsistency of feature rankings each time. The markers used in dpGLAS were only single random representatives among the co-segregating markers. But the genomic boundaries of these co-segregating members will guide to the genes of interest and the interpretation becomes easy for those markers that have very few co-segregating or single locus,

The dpGLAS we proposed here is specific for bi-parental population data with genic markers (**Fig 7**). We integrated bi-parental population data based on the one parent genotype that carried the genes of interest trait. This integration was proven to work since the dpGLAS model trained from 0.9 portion of the integration sample set was able to predict new samples originally from two independent sets. We assume that the gene effect on FHB resistance carried from one parent was gain-of-function, i.e., the other parent did not carry this functional gene. The marker genes having identical parental genotypes were unlikely of FHB resistant sites, and therefore were classified as a non-interest parent type (BrcBa). This idea will allow us to integrate more bi-parental or multi-parental population data together. It is worth noting that there were not many markers that were uncertain in dpGLAS design and majority markers were rationally excluded from the candidate pool.

**Fig 7.**
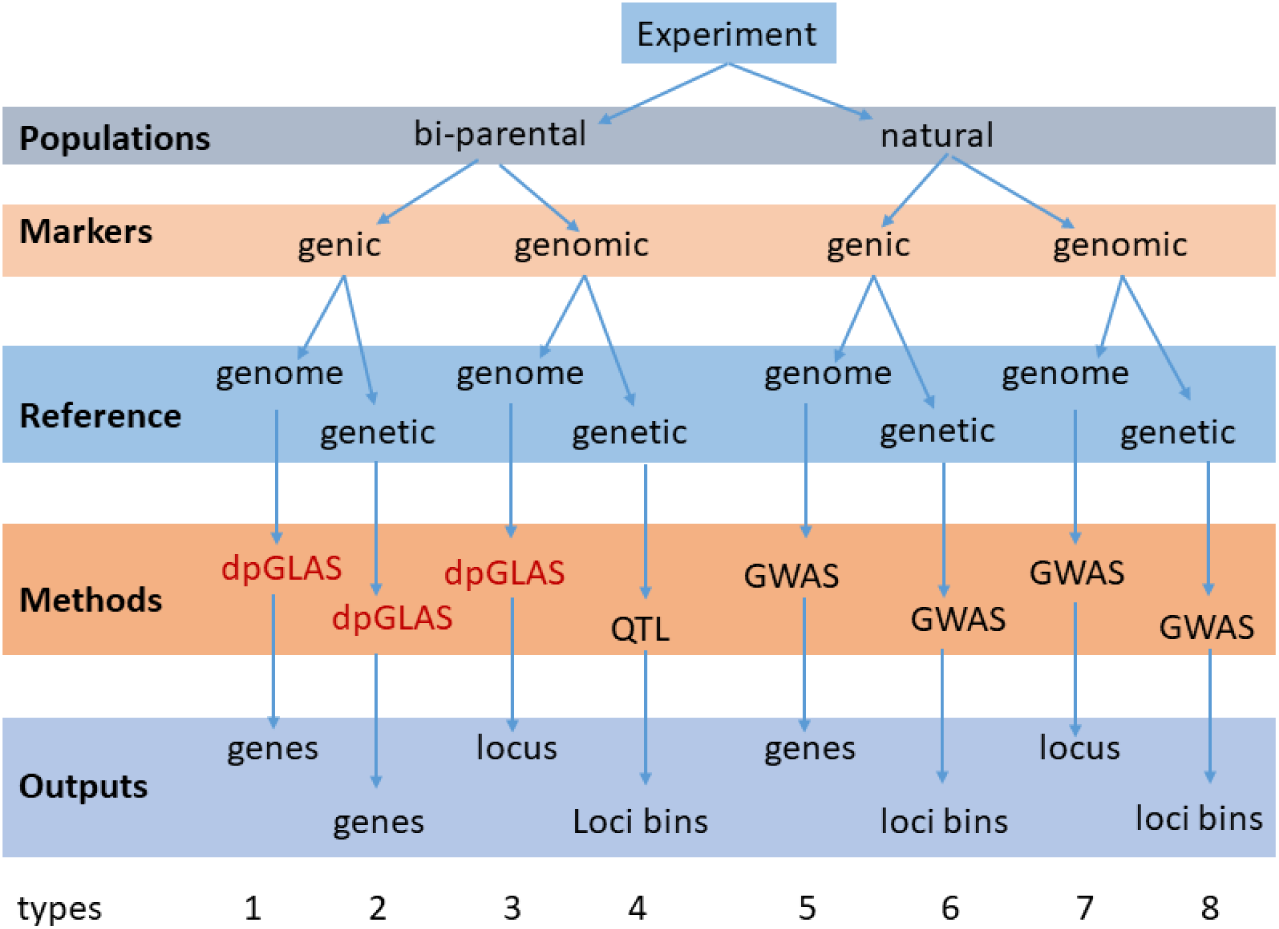
Six scenarios (bi-parental, natural, genic, genomic, genome, genetic) of experiments by three types of analytic methods (dpGLAS, QTL linkage, and GWAS). The output of these scenarios are individual genes, individual variation loci, or loci bin with certain cM range.

The genetic control of FHB resistance in both populations Br and cB was confirmed by their heritability[39]. Given that FHB resistance is governed by multigene and environment factor interactions[42], the SHAP score of each gene marker was calculated by taking the combination of many gene markers that have contributed to the prediction outcome. However, many gene markers that played a similar weight of effects may not have the same order in SHAP scores in different validation tests except a few top genes. Therefore we chose the markers that were in top list with high frequency in repeated validations. The convolving process of CNN is considering interactions among neighboring SNPs within different ranges of the kernel window, capturing the local epistasis effect. The gene marker order resulting from final CNN helped build the interaction network of multigene in FHB resistance. The top 10 markers of the bi-populations that had the common FHB resistance AC Barrie parent manifested none overlap. This reflected the complication of multiple genes and factors for the quantitative trait of FHB resistance. Also, the different types of bi-parental populations (DH and RIL) generated different segregation patterns which led marker existence in one set but loss in the other set because of monomorphic. The different variant sites detected from the two populations (Br, cB, BrcBa) would add up more FHB resistance associated variant sites. When we expanded the top list to 20 marker genes, overlap appeared in different sets, for example the PP2C gene. A different method, the conventional QTL mapping, applied to the same data increased a piece of support to our results. Five out of the 10 markers of the Br and 3 of 10 of the cB data sets were also detected by QTL mapping.

There are three scenarios of interpretations of the genic markers: i) a marker of the causal gene; ii) a marker of a big segment that harbored the causal gene from one parent; iii) both parents had the same gene but causative mutation at the marker site exerted gain-of-function of FHB resistance in one parent gene. dpGLAS was able to catch the genes in case i) and iii). When one marker gene together with many other genes passed into one single segment through co-segregating in one bi-parent population, the gene marker membership size could be narrowed down after being compared to another bi-parent population during data integration because the same gene marker may have different co-segregating in different bi-parental populations. Here we used the same bi-parental experiment data[39] and discovered 30 candidate genes that may be associated with FHB resistance in AC Barrie. These marker associated genes showed that the AC Barrie may confer resistance to FHB via a ROS and ABA central path. The activation of phosphoinositide-specific phospholipase C (PI-PLC) is one of the earliest responses triggered by the recognition of several microbe-associated molecular patterns (MAMPs) in plants and PI-PLC involves early ROS-regulated processes[43]. Alkaline/neutral invertase might act as a negative regulator in wheat disease resistance to Pst by balancing the ROS production[44]. Ascorbate peroxidase (APX) enzymes play a key role catalyzing the conversion of H_2_O_2_ into H_2_O [45]. In plant cells, increased glucose-6-phosphate dehydrogenase expression has been related to resistance to oxidative stress[46]. In the plant cell, ROS can directly cause strengthening of host cell walls, and ROS are also important signals mediating defense gene activation. Additionally, ROS mediates the establishment of systemic defenses (systemic acquired resistance [SAR]) through association with SA[47]. Rar1 is involved in wheat defense against stripe rust through SA to influence ROS accumulation and HR[48]. TPR likely acts downstream in the ABA signal transduction pathway[49]. Cullin-RING ubiquitin ligases regulates salicylic acid (SA) and ABA in plant immune signaling[50]. Ubiquitylation has been shown to play important roles in abscisic acid (ABA) signaling[51]. PP2Cs are vital phosphatases that play important roles in abscisic acid (ABA) signaling[52]. RPM1 is a plant immune receptor that specially recognizes pathogen-released effectors to activate effector-triggered immunity (ETI). RPM1 triggers ETI and hypersensitive response (HR) for disease resistance[53]. In a different bi-parental population with another FHB resistant parent AAC Tenacious, the ROS related protein families (hydroxylase, peroxidase, and oxidoreductase) were significantly enriched in the differential response genes (DrGs) to FHB inoculation in RNA-eq transcriptome data. We also located a very important candidate TraesCS2D02G080000 (encoding Ascorbate peroxidase, APX) discovered in cB population by dpGLAS within peak position of the most important QTL of AAC Tenacious and very close (< 0.3 mb) to tightly linked marker *Ppd-D1* [41]. This finding is important because Roblin and AAC Tenacious, both are monomorphic at *Ppd-D1* locus and it seems impossible to identify a gene/qtl in this region using a population derived from these two cultivars. Even this gene was not discovered using conventional differential gene expression analysis using same RNASeq data set.

In summary, we have developed a deep learning Genome-wide Linkage Association Study (dpGLAS) which differs from the conventional GWAS and QTL linkage mappings. We used dpGLAS to discover the FHB resistant genes from two bi-parental populations and fine mapped the top 30 gene candidates in the Canadian native wheat cv. AC Barrie by two independent sets and one integrated set analyses. Eight marker genes were supported by overlaps with conventional QTL mapping. Most of these genes were found associated with ABA-ROS central path. In RNA-seq validation data of another resistant wheat cv. AAC Tenacious with FHB inoculation, these ABA-ROS related protein families were significantly enriched in the differential response genes DrGs to FHB inoculation compared to susceptible Roblin. dpGLAS also discovered a common marker site in the different FHB bi-parental populations AC Barrie and AAC Tenacious, the TraesCS2D02G080000 encoding Ascorbate peroxidase (APX) which is very close (< 0.3 mb) to tightly linked an important marker *Ppd-D1*. This is the first report for fine mapping to gene levels using bi-parental population data. This discovery will help GS and GE in wheat breeding program.

## Materials and methods

### Plant population genotypes and traits

Marker data on two wheat bi-parental populations generated in a previous study by Thambugala et al (2020) [39] were used in this study. The first was a recombinant inbred line (RIL) population of 195 lines from the cross Cutler/AC Barrie (cB). AC Barrie is a hard red spring wheat that possesses an intermediate level of FHB resistance, while Cutler is a hard red spring wheat that is susceptible to FHB. The second population was a doubled haploid (DH) population of 184 lines developed from the cross AC Barrie/Reeder (Br), where Reeder (PI-613586) is a moderately susceptible hard red spring wheat. A total of 81,277 SNPs were mapped in the cB population and 81,587 in Br, with the Illumina Infinium iSelect 90K wheat SNP BeadChip[14] (**Table 1**). Each population was independently tested for FHB resistance in multiple field trials, which were randomized as alpha lattice designs. FHB incidence and severity data were collected from inoculated FHB nurseries and FHB visual rating index [VRI = (FHB incidence × FHB severity)/100] was calculated. Best linear unbiased predictors (BLUPs) were calculated. The heritability estimate made in the original study[39] was 0.84 for VRI in the cB RIL population and 0.89 for VRI in the Br DH population. For method validation, a different dataset used a DH population reported by Dhariwal et al (2020) [41] with a different FHB resistant parent AAC Tenacious than AC Barrie. AAC Innova was the FHB-susceptible parent. This dataset contained a total of 188 DH lines and their parents (AAC Innova and AAC Tenacious). A total 81,588 variant sites were evaluated by the wheat 90K Infinium iSelect SNP assay and the FHB visual rating index were used in the dpGLAS for validation purposes.

### RNA-seq data and analysis

RNA-seq data from AAC Tenacious and Roblin generated by Kirby et al (2020)[54] were downloaded from NCBI. AAC Tenacious is the only spring wheat cultivar in Canada to receive a resistant rating to FHB, and belongs to the Canada Prairie Spring Red market class. Roblin is a spring wheat cultivar that is susceptible to FHB. The sample information and NCBI accession numbers were listed (Supplemental **Table S5)**. The fastq reads from AAC Tenacious were mapped on the Wheat IWGSC refseqv1.0 by program STAR (version 2.7.9a)[55] with the option–geneCount. The bam files were used in IGV [56] for variant site inspection. The raw count data output from STAR were further analyzed by R package edgeR (version 3.34.0) for differential gene expression analysis[57]. Briefly, the biological coefficient of variation (BCV) values were estimated by a negative binomial model. The fold-change and significance were tested by the generalized linear model (GLM) and quasi-likelihood (QL) dispersion estimation testing (glmQLFTest). The differential FHB response genes (DrGs) were determined by comparing the difference between the ratios of inoculated over control of AAC Tenacious (Tfc) and inoculated over control of Roblin (Rfc) normalized by the absolute value of the ratio of the AAC Tenacious control and Roblin control (TRc) and a factor of approaching log (1.0):

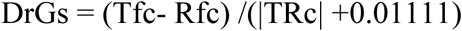

### Data integration and dpGLAS analytic design

These two types of parental populations, RIL and DH, are similar in that their recombinant progenies were all homozygous so that there were only two genotypes at any given locus, either from FHB resistant parent B (AC Barrie), or susceptible parent A (either Cutler or Reeder). The data integration was based on the parental information and the fact that these two populations had one parent in common, AC Barrie (B). The following three sets of integration were generated. A set labelled BrcB contained only consistent genotype markers and the markers with the same two parent genotypes in one of the two dataset were removed (Supplemental **Table S7)**. A set labelled BrcBa included and converted those marker genotypes with the same parent genotypes into A in the set where they reside in. A set labelled BrcBb included and converted those marker genotypes with the same parent genotypes into B in the set where they reside in. Comparison of these three integrated datasets helped optimization of dataset integration.

The dpGLAS is based on the assumption that the common parent (AC Barrie, B) in different populations contributed the same genes associated with the trait of interest and the markers to be used are anchored coding genes for the trait. The large non-coding regions in the genome, which are often the interval regions between markers, are not of interest to dpGLAS. The co-segregating sites in the population shared memberships. The first step is genotyping of parents, similar to traditional QTL mapping, and the second step is an association study. As shown in **Fig 1A**, the neural network deep learning dpGLAS included a feature selection by Dense Neural Network (DNN) and marker spatial effect by convolutional neural network (CNN).

### NN model and optimization

Neural network (NN) deep learning is an approach that repeatedly learns or adjusts the weight of each input feature (genotype marker) so that the sum of all features’ weights will explain the output (traits) at an error approaching zero (accuracy)[58].

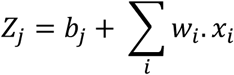

Where *j* is the neuron, *i* is the input feature, *Z*_*j*_ the output of a neuron, *b*_*j*_ the bias of the neuron, *w*_*i*_ the weight of a feature, and *x*_*i*_ the input feature value. The neurons in each NN layer will calculate their *w*_*i*_. During the learning process, two functions are implemented. The activation function transforms the product (*w*_*i*_.*x*_*i*_) to a mathematical pattern of value and the loss function monitors the progress in order to update the new weight. This radical idea was implemented in a variety of NN types and architectures.

The following considerations were taken into account for our study: the type of NN, NN architecture, and function parameters. The fully connected DNN evaluates the weight of each input feature by all neurons in a layer and reaches a result with least error. The CNN uses a sliding filter to test the combination of neighbour features together and the convoluted values from the sliding filter were summed which can infer feature interaction or epistasis. The CNN architecture included CNN and DNN modules. We tested the following CNN architectures. CNN11 was one Convolutional (Conv) layer and one Dense layer (**Supplemental Table S1**). CNN21 had two continuous Conv and one Dense where the input features were treated at first Conv and then the second Conv in one stream. CNN22 had two stream Conv and one Dense where the input features passed to two Conv streams independently. The Tensorflow v2.2.0 Python library package[59] was used in the development of the DNN and CNN models. The initializer, optimizer, activation, filter, pooling, batch, and epoch were also optimized by using the “GridSearchCV” function from the sklearn package[60] (Supplemental **Table S2**).

### SHAP score for marker ranking

The contribution of each individual feature (i.e. genotype marker) relative to all other features was examined by SHAP (SHapley Additive exPlanations) scores[61]. Features with large absolute Shapley values are considered important.

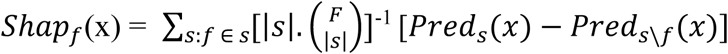

The SHAP score, *Shap*_*f*_(x), of feature *f* is the difference of scores between the whole feature set, *Pred*_*s*_(*x*), and the whole set without this feature (*Pred*_*s*\*f*_(*x*)), after a permutation 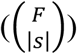 of the total number of features F was applied by the cardinality of each feature set combination, |*s*|. The Shap python library[61] was used for the SHAP score calculation and plot. The whole experiment was conducted as following steps: generate genotype input file, setup python sklearn package, run dpGLAS.py program with one input file, output 100 ranking lists of markers.

### DNN feature selection and validation

For each dataset, the fully connected DNN model was applied to evaluate all features. The top 100 features were used for 10-fold validation and repeated 100 times (**Fig 1B**). Each time a 0.1 portion of random samples were used as the test set, and the remaining 0.9 portion as the training set. The output of the predicted trait values of the test set by the trained model were compared to the test’s original trait values using Pearson correlation.

### CNN for epistasis markers and validation

The CNN was used to examine the spatial epistasis effect among marker genes. The CNN (CNN22) was applied to the top 100 features selected by the above DNN (**Fig 1B**). The CNN was run on the whole sample set 100 times, and each time the marker features were re-ordered based on their SHAP scores of previous run. The scores of each run were summed and the feature order with the highest sum score was updated and kept as the best marker order list of spatial effect. This list of spatial effect was further applied with 100 times of 10-fold validations (**Fig 1C**).

### Target annotation

For each of the bi-parental population datasets (Br, cB, BrcB, BrcBa, BrcBb, Tenacious), we selected the top 10 gene markers that appeared most frequently in the 100 10-fold validation in DNN. These 10 gene markers from five datasets by DNN were merged with the top 10 markers that were the most frequent markers in the 100 10-fold validation in CNN for final marker ranking. The memberships of the co-segregating markers were extracted (**Fig 1D**). The genomic loci of these markers were identified on th Chinese Spring wheat reference genome (IWGSC Refseq v1.0) by searching the T3/wheat database (https://triticeaetoolbox.org/wheat/). The genomic loci of these markers were used to resolve the uncertain locations of the paralogous genes or homologous genes on this hexaploid genome. Gene annotation was first conducted by blastx to the NCBI non redundant protein database and the genes were further analyzed by gene ontology and KEGG database search (https://www.genome.jp/kegg/genome/plant.html).

### Comparison with other methods

These two datasets (Br and cB) were analysed by the traditional QTL mapping. The genotype and phenotype data were input into software QTL IciMapping Version 4.2[62] for Inclusive Composite Interval Mapping for additive QTL (ICIM-ADD) analysis. The parameters of step of 0.1cM, probability in stepwise Regression (PIN 0.001), LOD of 2.5, and mapping function Haldane were used. The QTL mapping was repeated 100 times on randomly picked subsets of 90% samples. The results of QTL mapping and dpGLAS were compared and evaluated.

## Terms and abbreviations

Genic SNP marker: a SNP residing in a gene transcript
Genotype marker: a marker reflects the parent genotype
Allele B marker: allele carried by AC Barrie
Allele A marker: allele carried by the other parents of the bi-parental populations (either Cutler or Reeder)
Marker sites: polymorphic genetic loci
Marker number: the marker sites detected in the whole population
dpGLAS: neural network deep learning genome-wide linkage association study
GWAS: genome-wide association study
QTL: quantitative trait locus
DNN: Dense neural network
CNN: Convolutional neural network
RIL: recombinant inbred line
DH: doubled haploid
Br: genotyped DH population data of AC Barrie/Reeder cross
cB: genotyped RIL population data of Cutler/AC Barrie cross
BrcB: integrated data with exclusion of two identical parent genotypes existing in either set
BrcBa: integrated data, genotypes classified as non-B when two identical parent genotypes existing in one set
BrcBb: integrated data, genotypes classified as B when two identical parent genotypes existing in one set

## Declarations

### Ethics approval and consent to participate

Not applicable

### Consent for publication

Not applicable

### Availability of data and materials

All relevant data are within the paper and its Supplementary Data files. All code for dpGLAS will be publicly available on the dpGLAS GitHub repository: https://github.com/dpGLAS.

### Authors’ contributions

WX conceived of the study, implemented the dpGLAS program, conducted the data analysis and wrote the manuscript. MH, RD, HR provided genotyping data. All authors provided critical feedback and helped shape the research, analysis and manuscript. All authors read and approved the final manuscript.

## References

1. Horton MW, Hancock AM, Huang YS, Toomajian C, Atwell S, Auton A, Muliyati NW, Platt A, Sperone FG, Vilhjalmsson BJ, et al: Genome-wide patterns of genetic variation in worldwide Arabidopsis thaliana accessions from the RegMap panel. Nat Genet 2012, 44:212–216.

2. Asimit J, Zeggini E: Rare variant association analysis methods for complex traits. Annu Rev Genet 2010, 44:293–308.

3. Gibson G: Rare and common variants: twenty arguments. Nat Rev Genet 2012, 13:135–145.

4. Tam V, Patel N, Turcotte M, Bosse Y, Pare G, Meyre D: Benefits and limitations of genome-wide association studies. Nat Rev Genet 2019, 20:467–484.

5. Saint Pierre A, Genin E: How important are rare variants in common disease? Brief Funct Genomics 2014, 13:353–361.

6. Kearsey MJ, Farquhar AG: QTL analysis in plants; where are we now? Heredity (Edinb) 1998, 80 (Pt 2):137–142.

7. Drinkwater NR, Gould MN: The long path from QTL to gene. PLoS Genet 2012, 8:e1002975.

8. International Wheat Genome Sequencing C, investigators IRp, Appels R, Eversole K, Feuillet C, Keller B, Rogers J, Stein N, investigators Iw-gap, Pozniak CJ, et al: Shifting the limits in wheat research and breeding using a fully annotated reference genome. Science 2018, 361.

9. Lin HX, Qian HR, Zhuang JY, Lu J, Min SK, Xiong ZM, Huang N, Zheng KL: RFLP mapping of QTLs for yield and related characters in rice (Oryza sativa L.). Theor Appl Genet 1996, 92:920–927.

10. Yin X, Stam P, Dourleijn CJ, Kropff MJ: AFLP mapping of quantitative trait loci for yield-determining physiological characters in spring barley. Theoretical and Applied Genetics 1999, 99:244–253.

11. Ashkani S, Rafii MY, Rahim HA, Latif MA: Mapping of the quantitative trait locus (QTL) conferring partial resistance to rice leaf blast disease. Biotechnol Lett 2013, 35:799–810.

12. Curtolo M, Cristofani-Yaly M, Gazaffi R, Takita MA, Figueira A, Machado MA: QTL mapping for fruit quality in Citrus using DArTseq markers. BMC Genomics 2017, 18:289.

13. Nielsen R, Paul JS, Albrechtsen A, Song YS: Genotype and SNP calling from next-generation sequencing data. Nat Rev Genet 2011, 12:443–451.

14. Wang S, Wong D, Forrest K, Allen A, Chao S, Huang BE, Maccaferri M, Salvi S, Milner SG, Cattivelli L, et al: Characterization of polyploid wheat genomic diversity using a high-density 90,000 single nucleotide polymorphism array. Plant Biotechnol J 2014, 12:787–796.

15. Yousri NA, Fakhro KA, Robay A, Rodriguez-Flores JL, Mohney RP, Zeriri H, Odeh T, Kader SA, Aldous EK, Thareja G, et al: Whole-exome sequencing identifies common and rare variant metabolic QTLs in a Middle Eastern population. Nat Commun 2018, 9:333.

16. Wang MH, Cordell HJ, Van Steen K: Statistical methods for genome-wide association studies. Semin Cancer Biol 2019, 55:53–60.

17. Li M, Liu Y, Tao Y, Xu C, Li X, Zhang X, Han Y, Yang X, Sun J, Li W, et al: Identification of genetic loci and candidate genes related to soybean flowering through genome wide association study. BMC Genomics 2019, 20:987.

18. Xiong H, Guo H, Zhou C, Guo X, Xie Y, Zhao L, Gu J, Zhao S, Ding Y, Liu L: A combined association mapping and t-test analysis of SNP loci and candidate genes involving in resistance to low nitrogen traits by a wheat mutant population. PLoS One 2019, 14:e0211492.

19. Buzkova P: Linear regression in genetic association studies. PLoS One 2013, 8:e56976.

20. Chu BB, Keys KL, German CA, Zhou H, Zhou JJ, Sobel EM, Sinsheimer JS, Lange K: Iterative hard thresholding in genome-wide association studies: Generalized linear models, prior weights, and double sparsity. Gigascience 2020, 9.

21. Xu Y, Li P, Yang Z, Xu C: Genetic mapping of quantitative trait loci in crops. The Crop Journal 2017, 5:175–184.

22. Haley CS, Knott SA: A simple regression method for mapping quantitative trait loci in line crosses using flanking markers. Heredity (Edinb) 1992, 69:315–324.

23. Kao CH, Zeng ZB, Teasdale RD: Multiple interval mapping for quantitative trait loci. Genetics 1999, 152:1203–1216.

24. Van Ooijen JW: LOD significance thresholds for QTL analysis in experimental populations of diploid species. Heredity (Edinb) 1999, 83 (Pt 5):613–624.

25. Zhang F, Xie D, Liang M, Xiong M: Functional Regression Models for Epistasis Analysis of Multiple Quantitative Traits. PLoS Genet 2016, 12:e1005965.

26. Bzdok D, Altman N, Krzywinski M: Statistics versus machine learning. Nat Methods 2018, 15:233–234.

27. Ma W, Qiu Z, Song J, Li J, Cheng Q, Zhai J, Ma C: A deep convolutional neural network approach for predicting phenotypes from genotypes. Planta 2018, 248:1307–1318.

28. Montesinos-Lopez A, Montesinos-Lopez OA, Gianola D, Crossa J, Hernandez-Suarez CM: Multi-environment Genomic Prediction of Plant Traits Using Deep Learners With Dense Architecture. G3 (Bethesda) 2018, 8:3813–3828.

29. Liu Y, Wang D, He F, Wang J, Joshi T, Xu D: Phenotype Prediction and Genome-Wide Association Study Using Deep Convolutional Neural Network of Soybean. Front Genet 2019, 10:1091.

30. Figueroa M, Hammond-Kosack KE, Solomon PS: A review of wheat diseases-a field perspective. Mol Plant Pathol 2018, 19:1523–1536.

31. Giancaspro A, Giove SL, Zito D, Blanco A, Gadaleta A: Mapping QTLs for Fusarium Head Blight Resistance in an Interspecific Wheat Population. Front Plant Sci 2016, 7:1381.

32. Waldron BL, Moreno-Sevilla B, Anderson JA, Stack RW, Frohberg RC: RFLP Mapping of QTL for Fusarium Head Blight Resistance in Wheat. Crop Science 1999, 39:805–811.

33. Anderson JA, Stack RW, Liu S, Waldron BL, Fjeld AD, Coyne C, Moreno-Sevilla B, Fetch JM, Song QJ, Cregan PB, Frohberg RC: DNA markers for Fusarium head blight resistance QTLs in two wheat populations. Theoretical and Applied Genetics 2001, 102:1164–1168.

34. Buerstmayr H, Steiner B, Hartl L, Griesser M, Angerer N, Lengauer D, Miedaner T, Schneider B, Lemmens M: Molecular mapping of QTLs for Fusarium head blight resistance in spring wheat. II. Resistance to fungal penetration and spread. Theor Appl Genet 2003, 107:503–508.

35. Buerstmayr H, Lemmens M, Hartl L, Doldi L, Steiner B, Stierschneider M, Ruckenbauer P: Molecular mapping of QTLs for Fusarium head blight resistance in spring wheat. I. Resistance to fungal spread (Type II resistance). Theor Appl Genet 2002, 104:84–91.

36. Liu S, Pumphrey M, Gill B, Trick H, Zhang J, Dolezel J, Chalhoub B, Anderson J: Toward positional cloning ofFhb1, a major QTL for Fusarium head blight resistance in wheat. Cereal Research Communications 2008, 36:195–201.

37. Rawat N, Pumphrey MO, Liu S, Zhang X, Tiwari VK, Ando K, Trick HN, Bockus WW, Akhunov E, Anderson JA, Gill BS: Wheat Fhb1 encodes a chimeric lectin with agglutinin domains and a pore-forming toxin-like domain conferring resistance to Fusarium head blight. Nat Genet 2016, 48:1576–1580.

38. Li G, Zhou J, Jia H, Gao Z, Fan M, Luo Y, Zhao P, Xue S, Li N, Yuan Y, et al: Mutation of a histidine-rich calcium-binding-protein gene in wheat confers resistance to Fusarium head blight. Nat Genet 2019, 51:1106–1112.

39. Thambugala D, Brule-Babel AL, Blackwell BA, Fedak G, Foster AJ, MacEachern D, Gilbert J, Henriquez MA, Martin RA, McCallum BD, et al: Genetic analyses of native Fusarium head blight resistance in two spring wheat populations identifies QTL near the B1, Ppd-D1, Rht-1, Vrn-1, Fhb1, Fhb2, and Fhb5 loci. Theor Appl Genet 2020, 133:2775–2796.

40. Alonge M, Shumate A, Puiu D, Zimin AV, Salzberg SL: Chromosome-Scale Assembly of the Bread Wheat Genome Reveals Thousands of Additional Gene Copies. Genetics 2020, 216:599–608.

41. Dhariwal R, Henriquez MA, Hiebert C, McCartney CA, Randhawa HS: Mapping of Major Fusarium Head Blight Resistance from Canadian Wheat cv. AAC Tenacious. Int J Mol Sci 2020, 21.

42. Buerstmayr M, Huber K, Heckmann J, Steiner B, Nelson JC, Buerstmayr H: Mapping of QTL for Fusarium head blight resistance and morphological and developmental traits in three backcross populations derived from Triticum dicoccum x Triticum durum. Theor Appl Genet 2012, 125:1751–1765.

43. D’Ambrosio JM, Couto D, Fabro G, Scuffi D, Lamattina L, Munnik T, Andersson MX, Alvarez ME, Zipfel C, Laxalt AM: Phospholipase C2 Affects MAMP-Triggered Immunity by Modulating ROS Production. Plant Physiol 2017, 175:970–981.

44. Liu J, Han L, Huai B, Zheng P, Chang Q, Guan T, Li D, Huang L, Kang Z: Down-regulation of a wheat alkaline/neutral invertase correlates with reduced host susceptibility to wheat stripe rust caused by Puccinia striiformis. J Exp Bot 2015, 66:7325–7338.

45. Caverzan A, Passaia G, Rosa SB, Ribeiro CW, Lazzarotto F, Margis-Pinheiro M: Plant responses to stresses: Role of ascorbate peroxidase in the antioxidant protection. Genet Mol Biol 2012, 35:1011–1019.

46. Couee I, Sulmon C, Gouesbet G, El Amrani A: Involvement of soluble sugars in reactive oxygen species balance and responses to oxidative stress in plants. J Exp Bot 2006, 57:449–459.

47. Torres MA, Jones JD, Dangl JL: Reactive oxygen species signaling in response to pathogens. Plant Physiol 2006, 141:373–378.

48. Wang X, Wang Y, Liu P, Ding Y, Mu X, Liu X, Wang X, Zhao M, Huai B, Huang L, Kang Z: TaRar1 Is Involved in Wheat Defense against Stripe Rust Pathogen Mediated by YrSu. Front Plant Sci 2017, 8:156.

49. Schapire AL, Valpuesta V, Botella MA: TPR Proteins in Plant Hormone Signaling. Plant Signal Behav 2006, 1:229–230.

50. Furniss JJ, Spoel SH: Cullin-RING ubiquitin ligases in salicylic acid-mediated plant immune signaling. Front Plant Sci 2015, 6:154.

51. Miricescu A, Goslin K, Graciet E: Ubiquitylation in plants: signaling hub for the integration of environmental signals. J Exp Bot 2018, 69:4511–4527.

52. Park SY, Fung P, Nishimura N, Jensen DR, Fujii H, Zhao Y, Lumba S, Santiago J, Rodrigues A, Chow TF, et al: Abscisic acid inhibits type 2C protein phosphatases via the PYR/PYL family of START proteins. Science 2009, 324:1068–1071.

53. Yuan X, Wang Z, Huang J, Xuan H, Gao Z: Phospholipidase Ddelta Negatively Regulates the Function of Resistance to Pseudomonas syringae pv. Maculicola 1 (RPM1). Front Plant Sci 2018, 9:1991.

54. Nilsen KT, Walkowiak S, Kumar S, Molina OI, Randhawa HS, Dhariwal R, Byrns B, Pozniak CJ, Henriquez MA: Histology and RNA Sequencing Provide Insights Into Fusarium Head Blight Resistance in AAC Tenacious. Front Plant Sci 2020, 11:570418.

55. Dobin A, Davis CA, Schlesinger F, Drenkow J, Zaleski C, Jha S, Batut P, Chaisson M, Gingeras TR: STAR: ultrafast universal RNA-seq aligner. Bioinformatics 2013, 29:15–21.

56. Robinson JT, Thorvaldsdottir H, Winckler W, Guttman M, Lander ES, Getz G, Mesirov JP: Integrative genomics viewer. Nat Biotechnol 2011, 29:24–26.

57. Robinson MD, McCarthy DJ, Smyth GK: edgeR: a Bioconductor package for differential expression analysis of digital gene expression data. Bioinformatics 2010, 26:139–140.

58. Sze V, Chen Y-H, Yang T-J, Emer JS: Efficient Processing of Deep Neural Networks: A Tutorial and Survey. Proceedings of the IEEE 2017, 105:2295–2329.

59. Abadi M, Barham P, Chen J, Chen Z, Davis A, Dean J, Devin M, Ghemawat S, Irving G, al e: TensorFlow: A system for large-scale machine learning. 12th USENIX Symposium on Operating Systems Design and Implementation (OSDI 16) 2016:265--283.

60. Pedregosa F, Varoquaux G, Gramfort A, Michel V, Thirion B, Grisel O, Blondel M, Prettenhofer P, Weiss R, al. e: Scikit-learn: Machine Learning in Python. The Journal of Machine Learning Research 2011, 12:2825–2830.

61. Lundberg S, Lee S: A Unified Approach to Interpreting Model Predictions. NIPS’17: Proceedings of the 31st International Conference on Neural Information Processing Systems 2017:4768–4777.

62. Meng. L, Li. H, Zhang. L, Wang. J: QTL IciMapping: Integrated software for genetic linkage map construction and quantitative trait locus mapping in bi-parental populations. The Crop Journal 2015, 3:269–283.

