## supplementary for "Deep Learning Genome-wide Linkage Association Study for Wheat Fusarium Head Blight Resistance Genes Discovery"

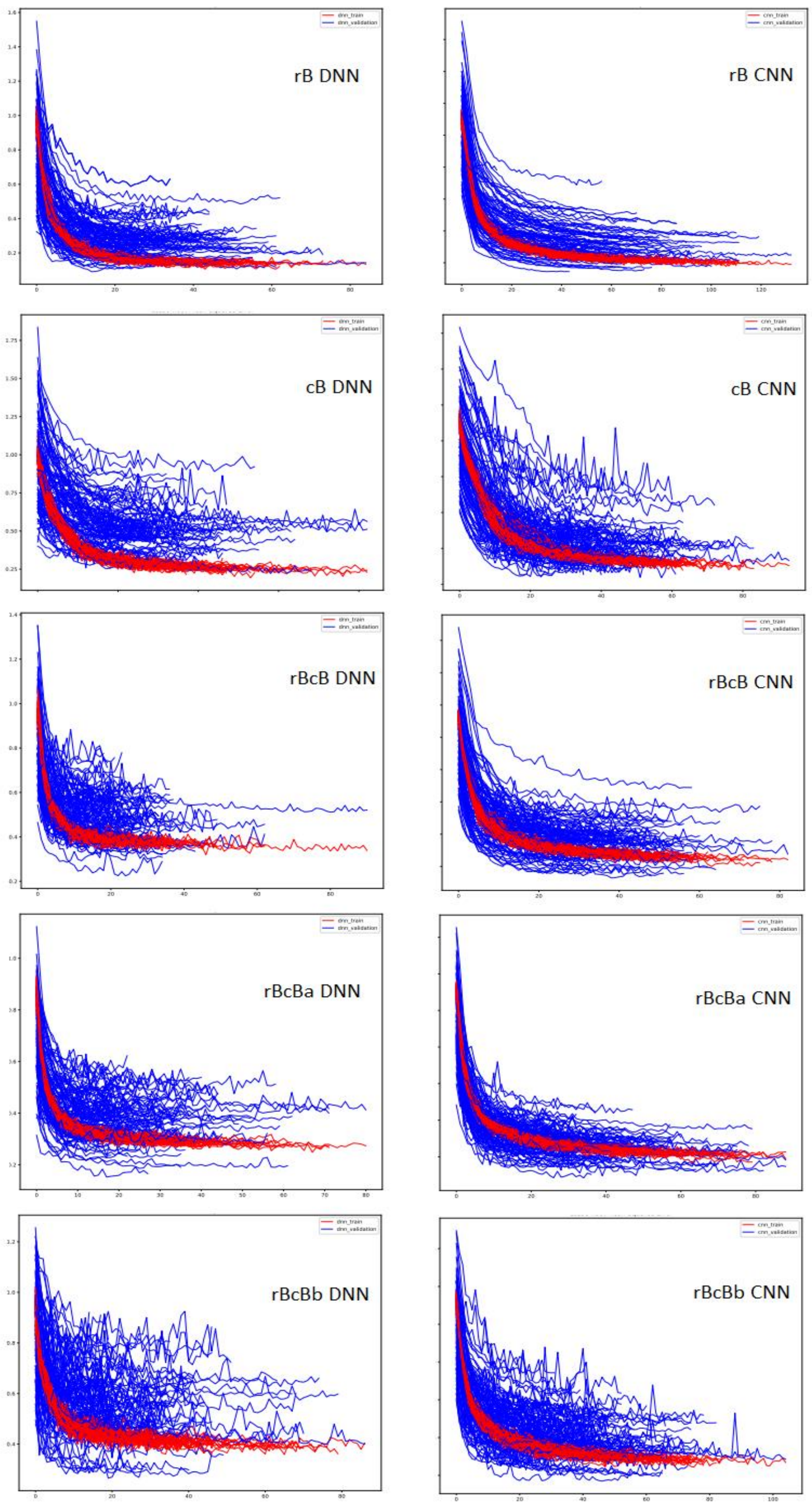


**Figure S1.** Loss of mean squared error (MSE) curves demonstrated the model performance of DNN and CNN in five data sets.

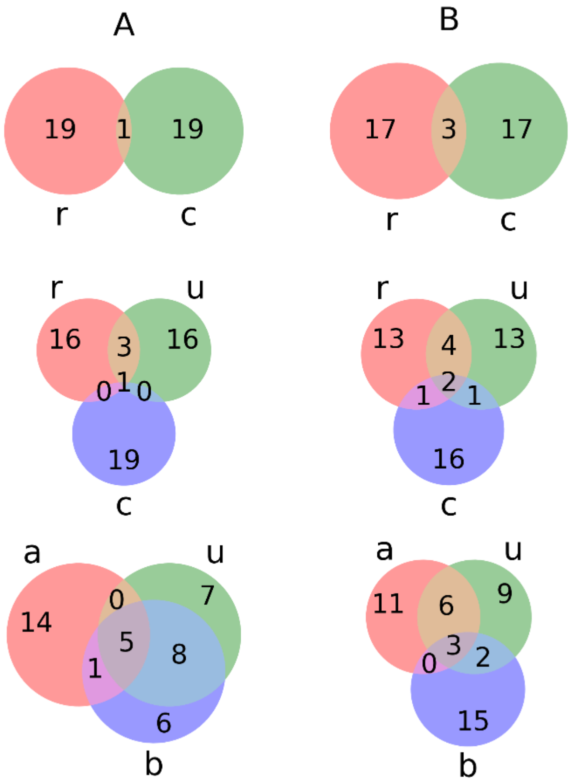


**Figure S2.** Overlap among the top 20 gene markers in the five datasets. A. gene markers from DNN validation. B. gene markers from CNN validation. Row 1: the two sets. Row 2: comparison of the two sets alone with the integrated set. Row 3: comparison of the three different integrations. One letter represents dataset: r: Br; c:cB; u: BrcB; a: BrcBa; b:BrcBb.

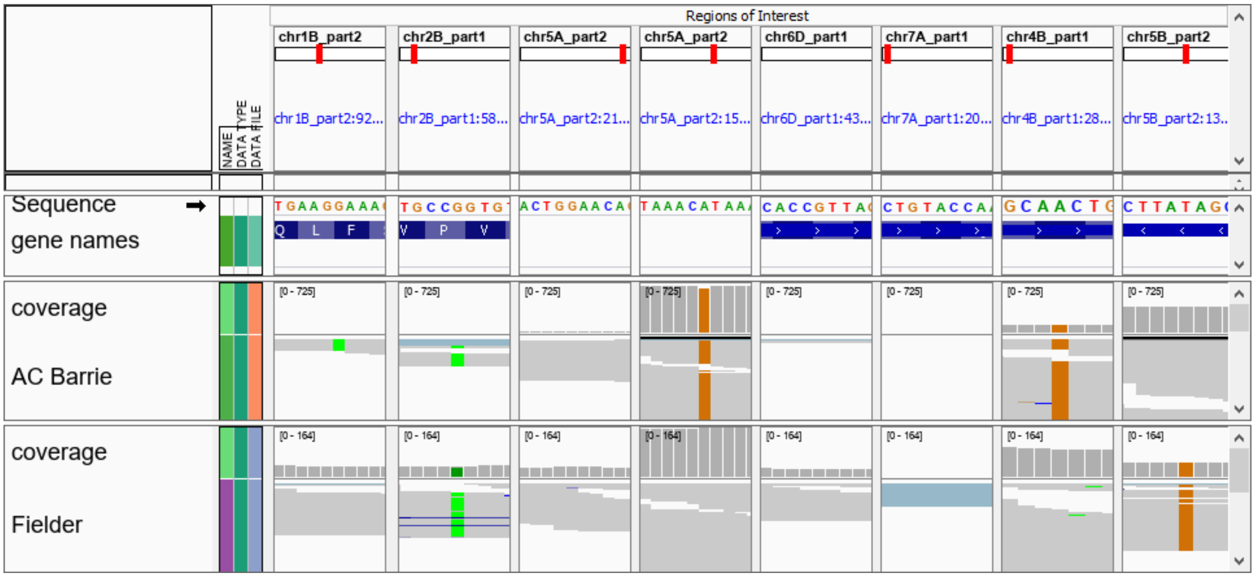


**Figure S3.** Eight marker associated genes and expression in AC Barrie and Fielder. The RNA-seq data were downloaded from NCBI SRA database and mapped on IWGSC RefSeq v1.0 by STAR program. Fielder was FHB susceptible wheat cultivar.

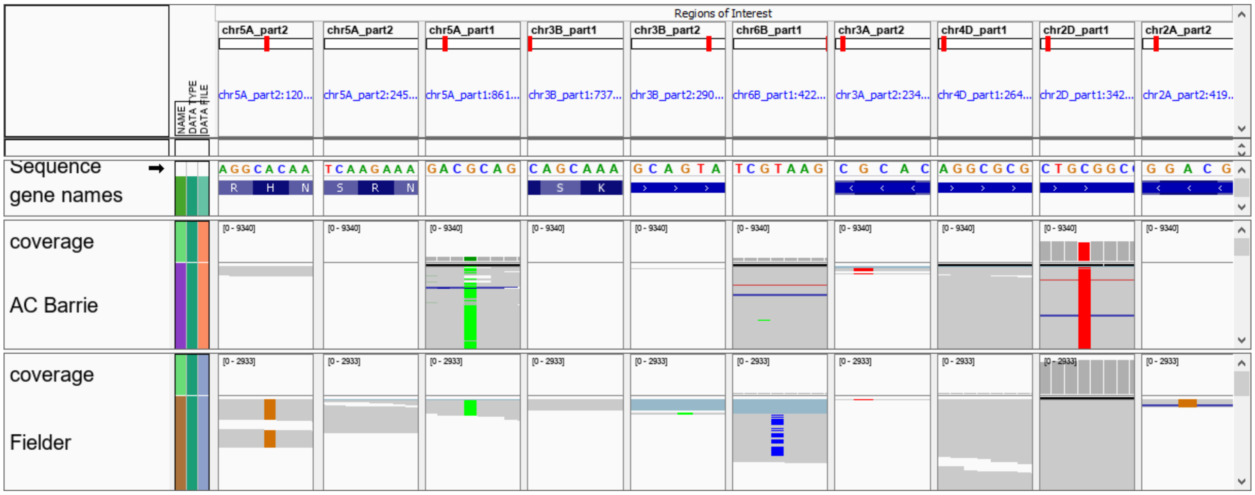


**Figure S4.** Ten variants of cB were in chr2A, 2D, 3A, 3B, 4D, and 5A. The RNA-seq data were downloaded from NCBI SRA database and mapped on IWGSC RefSeq v1.0 by STAR program. Fielder was FHB susceptible wheat cultivar.

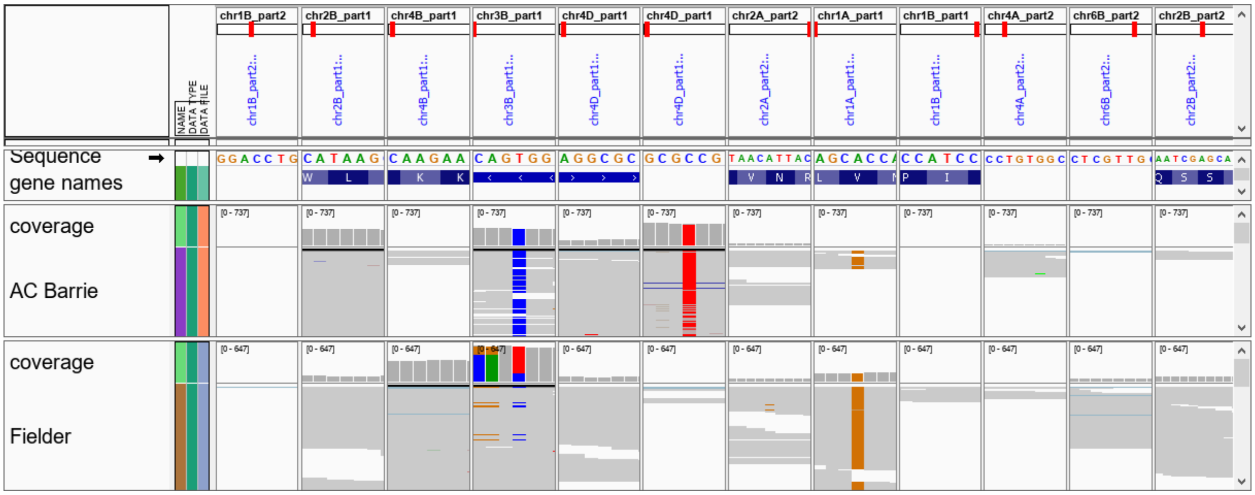


**Figure S5.** BrcBa, 12 variants/groups were found in chr1A,1B, 2A, 2B, 3B, 4A, 4B, 4D, and 6B

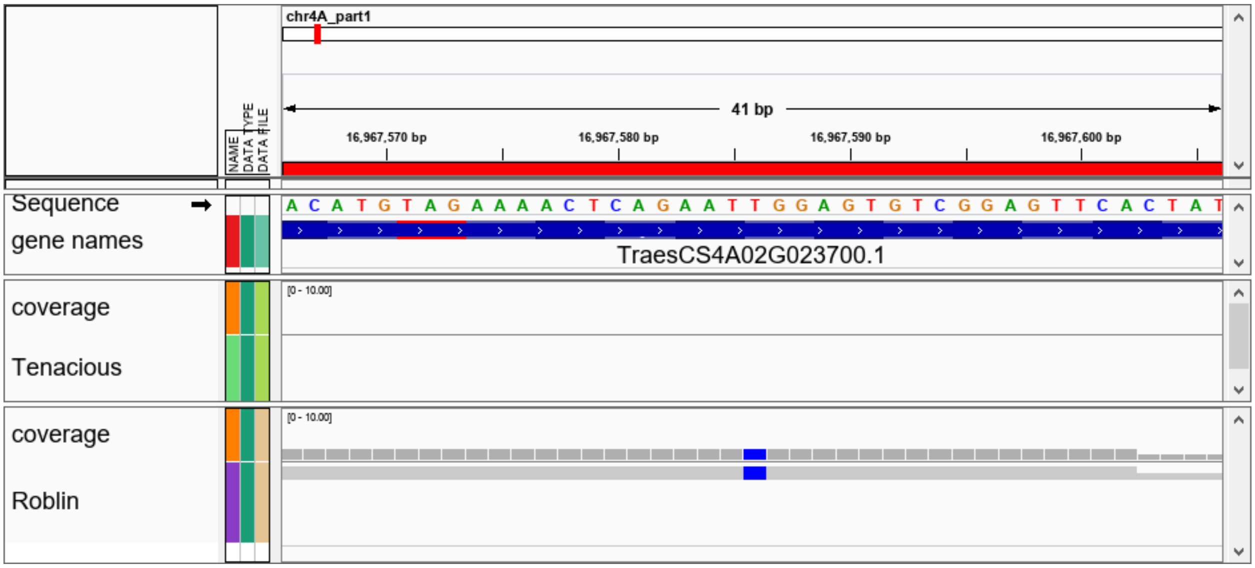


**Figure S6.** Gene TraesCS4A02G023700 coding for myosin-binding protein 3-like had a variant Ex_c7626_392 in chr4A:16967586. This is a missense variant, p.Leu663Ser. Both were under FHB inoculations.

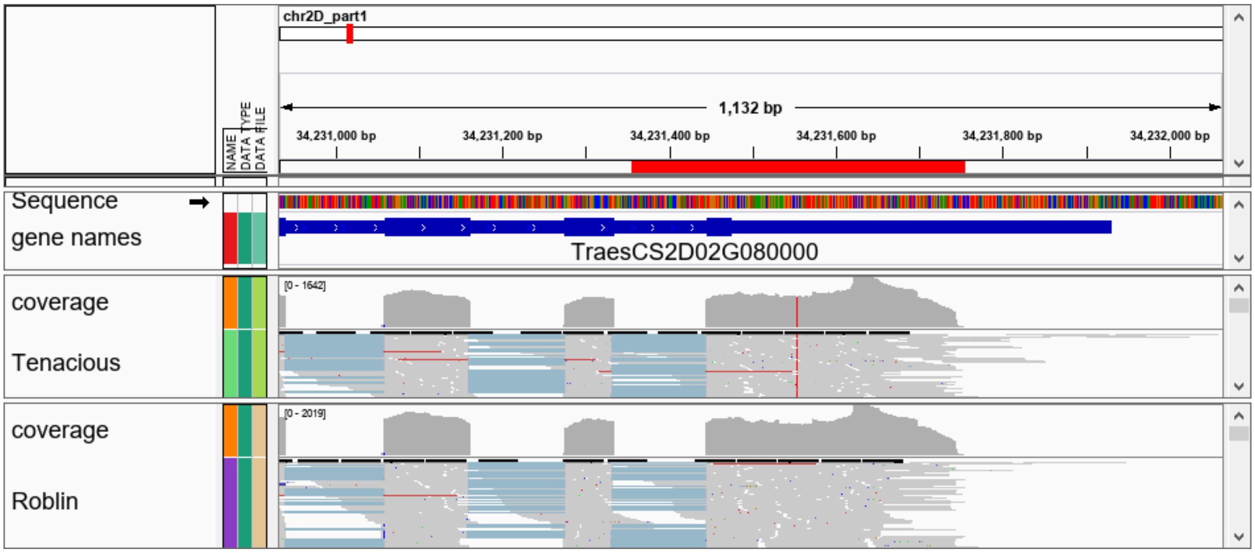


**Figure S7.** L-ascorbate peroxidase 2 (APX2) had a variant in intron. Both were under FHB inoculations.

.

**Table 1. NN architecture optimization**

| run | DNN | CNN11 | CNN12 | CNN22 |
|---|---|---|---|---|
| 1 | 0.546 | 0.382 | 0.473 | 0.526 |
| 2 | 0.299 | 0.249 | 0.201 | 0.456 |
| 3 | 0.506 | 0.584 | 0.430 | 0.515 |
| 4 | 0.415 | 0.406 | 0.627 | 0.567 |
| 5 | 0.456 | 0.233 | 0.530 | 0.497 |
| 6 | 0.327 | 0.294 | 0.422 | 0.424 |
| 7 | 0.309 | 0.152 | 0.421 | 0.344 |
| 8 | 0.575 | 0.567 | 0.645 | 0.783 |
| 9 | 0.637 | 0.591 | 0.531 | 0.655 |
| 10 | 0.128 | 0.062 | 0.188 | 0.266 |
| **mean** | **0.420** | **0.352** | **0.447** | **0.503** |
| SD | 0.147 | 0.177 | 0.147 | 0.140 |

DNN: dense
CNN11: one stream, 1 conv, 1 flatten dense
CNN12: one stream, 2 conv, 1 flatten dense
CNN22: two streams, 2 conv, 1 flatten dense

Table S2. NN parameters optimized

| run | 1 | 2 | 3 |
|---|---|---|---|
| **initializer** | TruncatedNormal | TruncatedNormal | TruncatedNormal |
| **optimizer** | RMSprop | RMSprop | Adamax |
| **activation** | tanh, linear | isru,linear | linear,linear |
| **filter_sz** | 10,4,4 | 6,10,4 | 20,4,4 |
| **pool_sz** | 6 | 2 | 3 |
| **initializer** | 'glorot_normal', 'he_normal','glorot_uniform','he_uniform','TruncatedNormal' | | |
| **optimizer** | 'Adam','Ftrl','SGD', 'RMSprop', 'Adagrad', 'Adadelta','Adamax', 'Nadam' | | |
| **activation** | activationss1 = ['tanh','linear','sigmoid',isru] | activationss2 = ['linear', isru] | |
| **filter_sz** | [6, 10, 20],[4, 10, 16],[4, 10, 16] | | |
| **pool_sz** | [2, 3, 4, 5, 6] | | |

Table S3. Candidate marker genes and co-segregating members in three data sets

| marker memberships | DNN | CNN | gene | site | annotation |
|---|---|---|---|---|---|
| **AC Barrie x Reeder (Br) (DH)** | | | | | |
| BobWhite_c16005_289 | 100 | 56 | TraesCS1D02G292600 | 1D391741516T/C | PLAC8 FAMILY PROTEIN |
| BS00010130_51 | | | | | |
| BS00010399_51 | | | | | |
| BS00011973_51 | | | | | |
| BS00012250_51 | | | | | |
| BS00064602_51 | | | | | |
| Ex_c102909_778 | | | | | |
| Excalibur_c23992_436 | | | | | |
| Excalibur_c48379_116 | | | | | |
| Excalibur_c48379_413 | | | | | |
| Excalibur_c48379_461 | | | | | |
| IACX11029 | | | | | |
| IACX11291 | | | | | |
| IACX5821 | | | | | |
| IACX9097 | | | | | |
| IACX9622 | | | | | |
| Kukri_c27693_260 | | | | | |
| Kukri_c27693_551 | | | | | |
| Kukri_rep_c104054_174 | | | | | |
| Ra_c62804_1641 | | | | | |
| RAC875_c7674_634 | | | | | |
| RAC875_c8888_802 | | | | | |
| RAC875_s110519_303 | | | | | |
| Tdurum_contig87938_342 | | | | | |
| wsnp_Ex_c23992_33235984 | | | | | |
| wsnp_JD_c11014_11565501 | | | | | |

| | | | | | |
|---|---|---|---|---|---|
| IACX1098 | 99 | 93 | TraesCS2B02G098500 | 2B58324675G/A | phosphoinositide phospholipase C 2-like |
| Kukri_c26288_419 | | | | | PWWP DOMAIN-CONTAINING PROTEIN (PTHR42851:SF7) Family: |
| Tdurum_contig54704_176 | | | | | |
| Kukri_c4221_1114 | 99 | 100 | | 5A672406766A/G | uncharacterized LOC119304065 |
| RAC875_rep_c106322_1091 | | | | | TPR_REGION DOMAIN-CONTAINING PROTEIN |
| Tdurum_contig29960_214 | | | | | |
| Tdurum_contig29960_91 | | | | | |
| BobWhite_c30500_527 | 93 | | TraesCS6D02G323900 | 6D430402119G/A | 7-deoxyloganetin glucosyltransferase-like |
| Excalibur_c23296_820 | | | | | |
| GENE-4566_348 | | | | | |
| Kukri_c25034_2042 | | | | | |
| Kukri_rep_c68504_1403 | | | | | |
| BS00022880_51 | 81 | | TraesCS7A02G046200 | 7A20969003A/G | disease resistance protein RPM1-like |
| CAP11_c1685_149 | 58 | 81 | | 5A607835124A/G | AP-1 complex subunit sigma-1 |
| Excalibur_c84439_196 | | | | | |
| Kukri_c6266_260 | | | | | |
| RAC875_c3964_752 | | | | | |
| tplb0050c03_1003 | | | | | |
| wsnp_Ex_c790_1554988 | | | | | |
| Ex_c101685_705 | 47 | 90 | TraesCS4B02G042300 | 4B28959979A/G | RAR1 gene |
| Ex_c101685_711 | | | | | |
| IACX557 | | | | | |
| wsnp_BF482960B_Ta_1_4 | | | | | |
| Tdurum_contig98569_290 | | 84 | TraesCS5B02G414800 | 5B589120307T/G | L-3-cyanoalanine synthase/cysteine synthase C1 |
| Tdurum_contig98569_56 | | | | | |
| **Cutler x AC Barrie (cB) (RIL)** | | | | | |
| BobWhite_rep_c49176_485 | 100 | | | 5A86181443G/A | polyubiquitin-like |
| BS00064387_51 | | | | | |
| Ex_c1344_1859 | | | | | |
| Excalibur_rep_c69282_651 | | | | | |
| IAAV4939 | | | | | |
| RFL_Contig4653_1217 | | | | | |
| Tdurum_contig81753_159 | | | | | |
| Tdurum_contig81753_70 | | | | | |
| wsnp_BE500291A_Ta_2_1 | | | | | |
| wsnp_Ex_c130_258776 | | | | | |
| wsnp_Ex_c130_259006 | | | | | |
| wsnp_Ex_c130_259533 | | | | | |
| wsnp_Ku_c16812_25759885 | | | | | |
| wsnp_Ku_c328_679106 | | | | | |
| wsnp_Ra_c2228_4310870 | | | | | |
| Excalibur_c11505_806 | 100 | 100 | TraesCS3B02G017800 | 3B7373756C/T | pentatricopeptide repeat-containing protein |
| RAC875_c30348_182 | | | | | |
| Tdurum_contig94629_258 | | | | | |
| tplb0059m03_622 | | | | | |
| BS00060666_51 | 98 | 99 | TraesCS3B02G493200 | 3B738752368G/A | homeobox protein LUMINIDEPENDENS-like |
| Tdurum_contig44343_1039 | 90 | 57 | TraesCS5A02G375500 | 5A573589851A/G | BTB/POZ domain-containing protein |
| Excalibur_c38128_422 | 83 | 68 | | 6B422702991T/C | |

| | | | | | |
|---|---|---|---|---|---|
| BS00062808_51 | 79 | 93 | TraesCS3A02G255900 | 3A477592042G/T | ABC transporter D family member 1-like |
| BobWhite_c8266_227 | 79 | 99 | TraesCS5A02G542600 | 5A698508163G/T | hexose carrier protein HEX6-like |
| RAC875_c8642_231 | | | | | |
| wsnp_Ex_rep_c107564_91144523 | 45 | 99 | TraesCS4D02G050600 | 4D26481498C/T | UDP-glucose 6-dehydrogenase 4-like |
| RAC875_c7319_195 | | 99 | TraesCS2D02G080000 | 2D34231554C/T | L-ascorbate peroxidase 2 |
| wsnp_CAP11_c3842_1829821 | | | | | |
| Tdurum_contig27861_306 | | 88 | TraesCS2A02G292900 | 2A504278926A/G | neutral/alkaline invertase 3 |
| **BrcBa set** | | | | | |
| wsnp_Ex_rep_c107564_91144523 | 100 | | TraesCS4D02G050600 | 4D26481498C/T | UDP-glucose 6-dehydrogenase 4-like |
| Tdurum_contig50326_645 | 100 | 100 | TraesCS3B02G000800 | 3B987585T/C | putative disease resistance protein RGA1 |
| RAC875_c19042_642 | 99 | 39 | TraesCS2A02G563500 | 2A764100959A/G | uncharacterized LOC119357316 |
| tplb0025b13_2054 | 98 | 66 | TraesCS1A02G005800 | 1A3387935A/G | protein MEI2-like 3 |
| IAAV3905 | 89 | 100 | TraesCS1B02G219100 | 1B396531147A/G | uncharacterized LOC119338372 |
| CAP12_c5519_132 | 88 | | | 4A521609833G/A | |
| Ex_c23792_486 | | | | | |
| Excalibur_c31814_298 | | | | | |
| IAAV1461 | | | | | |
| IAAV6223 | | | | | |
| IAAV7636 | | | | | |
| IAAV7786 | | | | | |
| Kukri_c13991_655 | | | | | |
| Kukri_c27037_112 | | | | | |
| Kukri_c74651_223 | | | | | |
| Kukri_c9578_2045 | | | | | |
| Ra_c30013_483 | | | | | |
| RAC875_c5394_1052 | | | | | |
| wsnp_BE398523A_Ta_2_1 | | | | | |
| wsnp_Ex_c11663_18779609 | | | | | |
| wsnp_Ex_c13623_21404172 | | | | | |
| wsnp_Ex_c2266_4247520 | | | | | |
| wsnp_Ex_c30876_39741201 | | | | | |
| wsnp_Ex_c3463_6348808 | | | | | |
| wsnp_Ex_c5492_9691241 | | | | | |
| wsnp_Ex_c5492_9691880 | | | | | |
| wsnp_Ex_c58286_59646499 | | | | | |
| wsnp_Ex_c5979_10480527 | | | | | |
| wsnp_Ex_c9464_15689857 | | | | | |
| wsnp_Ex_rep_c101826_87124211 | | | | | |
| wsnp_Ex_rep_c67779_66463916 | | | | | |
| wsnp_Ex_rep_c70327_69270561 | | | | | |
| wsnp_Ku_c10224_16965872 | | | | | |
| wsnp_Ku_c48043_54334230 | | | | | |
| wsnp_Ku_c5979_10559245 | | | | | |
| Excalibur_c31379_71 | 86 | | | 6B653212354G/A | uncharacterized LOC119323418 |

| | | | | | |
|---|---|---|---|---|---|
| BS00064602_51 | 40 | 98 | | 1B539930159A/G | |
| RAC875_c6922_291 | | 100 | TraesCS4B02G032600 | 4B24398838G/T | uncharacterized LOC119276596 |
| CAP8_rep_c5023_658 | | 100 | | 4D20579698C/T | uncharacterized LOC109778841 |
| wsnp_CAP11_c356_280910 | | | | | |
| RAC875_c2092_1020 | | 98 | TraesCS2B02G449200 | 2B641941825G/A | probable protein phosphatase 2C 44 |
| Kukri_c26288_419 | | 89 | TraesCS2B02G096600 | 2B56784859A/G | uncharacterized LOC119362187 |

Table S4. The top markers and associated genes from AAC Tenacious x Roblin set detected by dpGLAS and overlapped by conventional QTL mapping.

| dpGLAS marker members | DNN | CNN | gene | Site (IWSGCv2.0) | annotation | QTL mapping** | Site (v2.0) | distance |
|---|---|---|---|---|---|---|---|---|
| **AAC Innova x AAC Tenacious (DH)** | | | | | | | | |
| wsnp_CAP11_c3842_1829821 | 82 | | TraesCS2D02G080000 | 2D36482398* | L-ascorbate peroxidase 2 | Ppd-D1 | 2D36205433 | -276965 |
| RAC875_c7319_195 | 49 | 87 | TraesCS2D02G080000 | 2D36482487 | L-ascorbate peroxidase 2 | epistasis | | |
| Ex_c7626_392 | 100 | | TraesCS4A02G023700 | 4A16857343* | myosin-binding protein 3-like | | | |
| Ex_c7626_444 | | | TraesCS4A02G023700 | 4A16857414 | myosin-binding protein 3-like | | | |
| Ku_c7594_1179 | | | TraesCS4A02G023700 | 4A16856843 | myosin-binding protein 3-like | | | |
| Excalibur_c13054_1564 | 72 | 98 | TraesCS4A02G402100 | | RGA4 | | | |
| RAC875_c35979_263 | | | TraesCS4A02G597500LC | 4A678390139 | | | | |
| Kukri_c23629_120 | 91 | | TraesCS4A02G402100 | 4A677884111* | RGA4 | | | |
| RFL_Contig4334_379 | | | TraesCS4A02G401700 | | CYSTATHIONINE BETA-SYNTHASE | | | |
| wsnp_Ex_c33327_41834973 | 92 | | TraesCS5D02G240800 | | microtubule-associated protein 6-like | | | |
| BS00023627_51 | 98 | 97 | | 6A51480141 | | | | |
| wsnp_JD_c2180_3000498 | | | TraesCS6A02G079000 | | reductase | | | |
| IAAV5530 | 59 | 94 | TraesCS7B02G018300 | | RGA5-like | | | |
| wsnp_CAP7_c44_26549 | | | TraesCS7B02G018600 | 7B17091302* | soluble starch synthase (SS1) | wsnp_CAP7_c44_26549 | 7B17091249 | -53 |

The top 10 markers that appeared at least 80 times in the 100 subsets learning of either DNN or CNN. The markers that do not show details of DNN and CNN learnings are co-segregating members. *start site of the snp probe, not the actual snp site.

Table S5. The most different response to FHB compared the resistance Tenacious and the susceptible Roblin. 218 genes.

| chr | start | gene | Rfc | Tfc | TRc | logCPM3 | TRf | DrG |
|---|---|---|---|---|---|---|---|---|
| chr3B | 726621717 | TraesCS3B02G478300 | 0.0 | 12.3 | 0.0 | 2.9 | 12.3 | 1110.6 |
| chr5A | 708071305 | TraesCS5A02G556900 | 0.0 | 10.6 | 0.0 | 1.2 | 10.6 | 956.8 |
| chr3D | 27054173 | TraesCS3D02G061800 | 0.0 | 10.6 | 0.0 | 1.2 | 10.6 | 955.3 |
| chr5A | 14958561 | TraesCS5A02G024900LC | 0.0 | 9.5 | 0.0 | 0.2 | 9.5 | 854.5 |
| chr7B | 732577242 | TraesCS7B02G795900LC | 0.0 | 9.4 | 0.0 | 0.1 | 9.4 | 849.4 |
| chr7B | 732577653 | TraesCS7B02G796000LC | 0.0 | 9.4 | 0.0 | 0.1 | 9.4 | 848.1 |
| chr6B | 562487341 | TraesCS6B02G314600 | 0.0 | 9.3 | 0.0 | 0.0 | 9.3 | 835.8 |
| chr7A | 13928557 | TraesCS7A02G037100LC | 0.0 | 8.9 | 0.0 | -0.4 | 8.9 | 799.1 |
| chr6B | 599700618 | TraesCS6B02G618700LC | 0.0 | 8.0 | 0.0 | -1.2 | 8.0 | 718.2 |
| chr5A | 591868022 | TraesCS5A02G397500 | 0.0 | 7.7 | 0.0 | -1.4 | 7.7 | 695.0 |
| chrUn | 51332408 | TraesCSU02G066200 | 0.0 | 7.5 | 0.0 | -1.6 | 7.5 | 673.7 |
| chr6B | 10934559 | TraesCS6B02G017100 | 0.0 | 7.0 | 0.0 | -1.9 | 7.0 | 631.9 |
| chr3A | 9555972 | TraesCS3A02G012400 | 5.1 | 12.0 | 0.0 | 2.6 | 6.8 | 616.4 |
| chr3A | 534381815 | TraesCS3A02G430100LC | 6.6 | 11.4 | 0.0 | 2.1 | 4.9 | 437.2 |
| chr1A | 469309603 | TraesCS1A02G275000 | -6.4 | -1.8 | 0.0 | -1.3 | 4.6 | 369.8 |
| chr7D | 42799929 | TraesCS7D02G072900 | 4.2 | 7.8 | 0.0 | -1.3 | 3.6 | 328.2 |
| chr6B | 279592487 | TraesCS6B02G211600 | 4.6 | 7.9 | 0.0 | -1.2 | 3.4 | 304.0 |
| chr4A | 604165624 | TraesCS4A02G470200LC | 4.1 | 7.5 | 0.0 | -1.5 | 3.4 | 303.3 |
| chr6A | 584997804 | TraesCS6A02G352900 | 5.6 | 8.9 | 0.0 | -0.3 | 3.3 | 296.0 |
| chr7A | 16790825 | TraesCS7A02G043600LC | 3.6 | 6.9 | 0.0 | -1.9 | 3.3 | 295.8 |
| chr7A | 12198019 | TraesCS7A02G030100 | -1.6 | 7.0 | 0.0 | -0.6 | 8.6 | 294.8 |
| chr4B | 24759437 | TraesCS4B02G033600 | 8.7 | 12.0 | 0.0 | 2.7 | 3.3 | 292.7 |
| chr1B | 669191911 | TraesCS1B02G452300 | 7.6 | 10.8 | 0.0 | 1.5 | 3.1 | 282.9 |
| chr7B | 2328395 | TraesCS7B02G004500 | 7.5 | 10.6 | 0.0 | 1.3 | 3.1 | 278.8 |
| chr1D | 35815390 | TraesCS1D02G064700LC | 5.6 | 8.7 | 0.0 | -0.5 | 3.1 | 277.4 |
| chr3B | 542722070 | TraesCS3B02G336500 | 6.6 | 9.7 | 0.0 | 0.5 | 3.1 | 276.7 |
| chr2B | 787532311 | TraesCS2B02G606400 | 4.9 | 7.9 | 0.0 | -1.2 | 3.1 | 275.3 |
| chr1B | 501176471 | TraesCS1B02G510800LC | 7.1 | 10.0 | 0.0 | 0.8 | 2.9 | 265.3 |
| chr2B | 745765055 | TraesCS2B02G549700 | 5.3 | 8.2 | 0.0 | -0.9 | 2.9 | 258.5 |
| chr5A | 590378540 | TraesCS5A02G544700LC | 4.9 | 7.7 | 0.0 | -1.4 | 2.8 | 252.0 |
| chr2D | 581569989 | TraesCS2D02G592300LC | 4.9 | 7.7 | 0.0 | -1.3 | 2.8 | 249.1 |
| chr2B | 48140846 | TraesCS2B02G102000LC | 4.2 | 6.9 | 0.0 | -1.9 | 2.8 | 249.0 |
| chr6B | 705628064 | TraesCS6B02G441300 | 1.1 | 8.8 | 0.0 | 1.0 | 7.7 | 246.9 |
| chr3B | 655027561 | TraesCS3B02G418000 | -6.3 | -2.0 | 0.0 | -1.4 | 4.2 | 246.9 |
| chr1B | 16670902 | TraesCS1B02G033100LC | 8.2 | 10.9 | 0.0 | 1.6 | 2.7 | 243.0 |
| chr4D | 456690657 | TraesCS4D02G286300 | 4.8 | 7.5 | 0.0 | -1.4 | 2.7 | 242.4 |
| chr7D | 549398311 | TraesCS7D02G584500LC | 4.9 | 7.6 | 0.0 | -1.4 | 2.7 | 241.8 |

| | | | | | | | | |
|---|---|---|---|---|---|---|---|---|
| chr3B | 666790281 | TraesCS3B02G428000 | 4.8 | 7.4 | 0.0 | -1.5 | 2.6 | 235.5 |
| chr7A | 66446673 | TraesCS7A02G109500 | 5.5 | 8.1 | 0.0 | -1.0 | 2.6 | 233.8 |
| chr1D | 334451082 | TraesCS1D02G243900 | -3.2 | 0.5 | 0.0 | -1.3 | 3.7 | 231.3 |
| chr6B | 39343093 | TraesCS6B02G079900LC | 5.1 | 7.7 | 0.0 | -1.3 | 2.5 | 228.9 |
| chr3A | 41484927 | TraesCS3A02G082700LC | 8.4 | 10.8 | 0.0 | 1.6 | 2.4 | 218.4 |
| chr6D | 457589885 | TraesCS6D02G372300 | 6.6 | 9.0 | 0.0 | -0.1 | 2.4 | 215.8 |
| chr2B | 649479177 | TraesCS2B02G454600 | 6.0 | 8.3 | 0.0 | -0.8 | 2.4 | 212.7 |
| chr5A | 565247984 | TraesCS5A02G364700 | 3.2 | 6.4 | 0.0 | -1.1 | 3.2 | 211.7 |
| chr1A | 535446433 | TraesCS1A02G351000 | 5.0 | 0.4 | 0.0 | 2.5 | -4.5 | -202.0 |
| chr1D | 7038659 | TraesCS1D02G014500 | 7.4 | 5.2 | 0.0 | -1.9 | -2.3 | -203.2 |
| chr3B | 690112426 | TraesCS3B02G449600 | 9.3 | 7.0 | 0.0 | -0.3 | -2.3 | -203.5 |
| chr2D | 12125609 | TraesCS2D02G029400 | 11.2 | 8.9 | 0.0 | 1.8 | -2.3 | -203.5 |
| chr5B | 672953006 | TraesCS5B02G507600 | 8.8 | 6.5 | 0.0 | -0.8 | -2.3 | -204.7 |
| chr5B | 616104269 | TraesCS5B02G443000 | 10.7 | 8.4 | 0.0 | 1.2 | -2.3 | -205.6 |
| chr5A | 559038447 | TraesCS5A02G356800 | 10.0 | 7.7 | 0.0 | 0.4 | -2.3 | -205.8 |
| chr2B | 631140937 | TraesCS2B02G668000LC | 8.7 | 6.4 | 0.0 | -0.8 | -2.3 | -206.3 |
| chr2A | 752998839 | TraesCS2A02G542000 | 13.6 | 11.3 | 0.0 | 4.3 | -2.3 | -206.3 |
| chr5A | 545835784 | TraesCS5A02G472500LC | 8.3 | 6.1 | 0.0 | -1.2 | -2.3 | -206.4 |
| chr2B | 560185632 | TraesCS2B02G395100 | 8.9 | 6.6 | 0.0 | -0.6 | -2.3 | -207.0 |
| chr5B | 616026070 | TraesCS5B02G442900 | 13.2 | 10.9 | 0.0 | 3.9 | -2.3 | -207.6 |
| chr4B | 660440304 | TraesCS4B02G589000LC | 9.0 | 6.7 | 0.0 | -0.5 | -2.3 | -207.7 |
| chr3B | 770116482 | TraesCS3B02G781500LC | 10.4 | 8.1 | 0.0 | 0.8 | -2.3 | -208.0 |
| chr5B | 616160236 | TraesCS5B02G443100 | 11.3 | 9.0 | 0.0 | 1.8 | -2.3 | -208.1 |
| chr7D | 160708608 | TraesCS7D02G202000 | 8.9 | 6.6 | 0.0 | -0.7 | -2.3 | -208.5 |
| chrUn | 299068480 | TraesCSU02G201300 | 8.3 | 6.0 | 0.0 | -1.2 | -2.3 | -208.6 |
| chr3D | 18699762 | TraesCS3D02G039000LC | 8.3 | 6.0 | 0.0 | -1.2 | -2.3 | -209.2 |
| chr5A | 35689851 | TraesCS5A02G038600 | 9.4 | 7.1 | 0.0 | -0.2 | -2.3 | -209.6 |
| chr4A | 709997217 | TraesCS4A02G441700 | 10.1 | 7.8 | 0.0 | 0.6 | -2.3 | -209.6 |
| chrUn | 335555235 | TraesCSU02G226400 | 15.7 | 13.4 | 0.0 | 6.5 | -2.3 | -210.5 |
| chr2D | 626601326 | TraesCS2D02G672100LC | 8.8 | 6.5 | 0.0 | -0.7 | -2.4 | -211.7 |
| chrUn | 391042971 | TraesCSU02G253900 | 11.5 | 9.1 | 0.0 | 2.1 | -2.4 | -212.0 |
| chr2A | 9785223 | TraesCS2A02G021200LC | 8.9 | 6.5 | 0.0 | -0.8 | -2.4 | -213.4 |
| chr7A | 76355063 | TraesCS7A02G118200 | 11.0 | 8.6 | 0.0 | 1.5 | -2.4 | -214.5 |
| chr7A | 732048458 | TraesCS7A02G800200LC | 7.9 | 5.5 | 0.0 | -1.5 | -2.4 | -214.8 |
| chr1B | 599499788 | TraesCS1B02G369300 | 7.8 | 5.4 | 0.0 | -1.6 | -2.4 | -216.2 |
| chr3B | 39451770 | TraesCS3B02G066000 | 8.7 | 6.3 | 0.0 | -0.8 | -2.4 | -216.9 |
| chr7A | 161995897 | TraesCS7A02G199200 | 12.1 | 9.7 | 0.0 | 2.7 | -2.4 | -218.0 |
| chr5B | 513325497 | TraesCS5B02G329200 | 9.6 | 7.2 | 0.0 | 0.1 | -2.4 | -218.7 |
| chr2D | 626598874 | TraesCS2D02G551100 | 8.7 | 6.3 | 0.0 | -0.9 | -2.4 | -218.7 |
| chr4A | 683800725 | TraesCS4A02G411200 | 7.6 | 5.2 | 0.0 | -1.8 | -2.4 | -220.1 |
| chr1B | 636768443 | TraesCS1B02G409400 | 8.4 | 6.0 | 0.0 | -1.2 | -2.4 | -220.2 |
| chrUn | 318560620 | TraesCSU02G215000 | 9.3 | 6.9 | 0.0 | -0.4 | -2.5 | -221.1 |
| chr7B | 89911529 | TraesCS7B02G080100 | 11.5 | 9.0 | 0.0 | 2.0 | -2.5 | -221.4 |
| chr4A | 688850929 | TraesCS4A02G614800LC | 10.6 | 8.1 | 0.0 | 1.0 | -2.5 | -221.5 |

| | | | | | | | | |
|---|---|---|---|---|---|---|---|---|
| chr2A | 775734221 | TraesCS2A02G584328 | 9.1 | 6.6 | 0.0 | -0.6 | -2.5 | -222.5 |
| chr5D | 420595530 | TraesCS5D02G327700 | 9.3 | 6.8 | 0.0 | -0.3 | -2.5 | -223.6 |
| chr5A | 622144844 | TraesCS5A02G596700LC | 8.1 | 5.6 | 0.0 | -1.4 | -2.5 | -224.2 |
| chr6D | 52128449 | TraesCS6D02G086500 | 7.8 | 5.3 | 0.0 | -1.6 | -2.5 | -224.6 |
| chr2D | 23336696 | TraesCS2D02G058800 | 10.7 | 8.2 | 0.0 | 1.1 | -2.5 | -225.6 |
| chr4D | 435011783 | TraesCS4D02G264500 | 7.8 | 5.3 | 0.0 | -1.7 | -2.5 | -226.0 |
| chr4A | 705544563 | TraesCS4A02G435300 | 14.0 | 11.5 | 0.0 | 4.7 | -2.5 | -227.1 |
| chr7D | 35856753 | TraesCS7D02G064800 | 8.3 | 5.8 | 0.0 | -1.2 | -2.5 | -228.3 |
| chr4D | 738967 | TraesCS4D02G001200 | 11.9 | 9.4 | 0.0 | 2.5 | -2.5 | -228.7 |
| chr6A | 584593648 | TraesCS6A02G351700 | 8.5 | 5.9 | 0.0 | -1.1 | -2.5 | -229.0 |
| chr5B | 615951078 | TraesCS5B02G442800 | 10.4 | 7.8 | 0.0 | 0.8 | -2.6 | -230.5 |
| chr7D | 90508723 | TraesCS7D02G141400 | 7.3 | 4.8 | 0.0 | -2.0 | -2.6 | -230.6 |
| chr3B | 30679098 | TraesCS3B02G059100 | 14.0 | 11.4 | 0.0 | 4.7 | -2.6 | -231.8 |
| chr4B | 562436091 | TraesCS4B02G279300 | 11.6 | 9.0 | 0.0 | 2.1 | -2.6 | -232.2 |
| chr7B | 498969118 | TraesCS7B02G469500LC | 7.4 | 4.8 | 0.0 | -2.0 | -2.6 | -233.4 |
| chr2B | 158863583 | TraesCS2B02G183900 | 7.7 | 5.1 | 0.0 | -1.7 | -2.6 | -233.7 |
| chrUn | 234833819 | TraesCSU02G290300LC | 9.9 | 7.3 | 0.0 | 0.3 | -2.6 | -234.4 |
| chr2D | 10432444 | TraesCS2D02G023100 | 7.8 | 0.4 | 0.0 | -0.5 | -7.4 | -236.0 |
| chrUn | 68268248 | TraesCSU02G123400LC | 9.6 | 7.0 | 0.0 | 0.0 | -2.6 | -236.8 |
| chr2A | 771609443 | TraesCS2A02G578000 | 15.9 | 13.2 | 0.0 | 6.6 | -2.6 | -237.1 |
| chr2D | 556350451 | TraesCS2D02G446000 | 9.3 | 6.6 | 0.0 | -0.3 | -2.7 | -238.7 |
| chr7A | 735763341 | TraesCS7A02G566400 | 8.3 | 5.7 | 0.0 | -1.2 | -2.7 | -239.4 |
| chr7A | 50136029 | TraesCS7A02G086000 | 11.0 | 8.4 | 0.0 | 1.5 | -2.7 | -240.0 |
| chr2B | 680662103 | TraesCS2B02G726500LC | 9.2 | 6.5 | 0.0 | -0.4 | -2.7 | -240.6 |
| chr1B | 465364821 | TraesCS1B02G264500 | 7.9 | 5.2 | 0.0 | -1.6 | -2.7 | -245.0 |
| chr6D | 422293144 | TraesCS6D02G313900 | 8.0 | 5.3 | 0.0 | -1.5 | -2.7 | -245.1 |
| chr1D | 8796980 | TraesCS1D02G020400 | 9.5 | 6.7 | 0.0 | -0.1 | -2.7 | -246.1 |
| chr7B | 525955673 | TraesCS7B02G491100LC | 8.2 | 5.4 | 0.0 | -1.4 | -2.8 | -247.7 |
| chr6D | 18533353 | TraesCS6D02G043900 | 10.0 | 7.2 | 0.0 | 0.3 | -2.8 | -249.6 |
| chr1A | 559881495 | TraesCS1A02G393300 | 9.2 | 6.4 | 0.0 | -0.4 | -2.8 | -251.6 |
| chr4B | 43716062 | TraesCS4B02G061800LC | 10.9 | 8.1 | 0.0 | 1.3 | -2.8 | -252.1 |
| chr1B | 146116375 | TraesCS1B02G207200LC | 6.2 | 0.4 | 0.0 | -1.9 | -5.8 | -252.2 |
| chr5D | 497957559 | TraesCS5D02G448200 | 8.3 | 5.5 | 0.0 | -1.3 | -2.8 | -252.5 |
| chr6D | 32318862 | TraesCS6D02G066000 | 7.9 | 5.1 | 0.0 | -1.6 | -2.8 | -254.5 |
| chr7D | 575408378 | TraesCS7D02G456700 | 7.5 | 4.6 | 0.0 | -1.9 | -2.8 | -255.7 |
| chr2A | 125384772 | TraesCS2A02G180200LC | 9.4 | 6.6 | 0.0 | -0.3 | -2.8 | -256.2 |
| chr5A | 480528810 | TraesCS5A02G270300 | 8.3 | 5.4 | 0.0 | -1.3 | -2.8 | -256.3 |
| chr2A | 682633748 | TraesCS2A02G429700 | 8.7 | 5.8 | 0.0 | -0.9 | -2.9 | -257.7 |
| chr1A | 337960 | TraesCS1A02G001600 | 2.7 | -0.8 | 0.0 | -1.1 | -3.5 | -258.3 |
| chrUn | 345125707 | TraesCSU02G233000 | 13.1 | 10.3 | 0.0 | 3.8 | -2.9 | -259.2 |
| chr5D | 288811090 | TraesCS5D02G258700LC | 10.5 | 7.6 | 0.0 | 0.9 | -2.9 | -259.9 |
| chr5B | 658808501 | TraesCS5B02G488500 | 9.2 | 6.3 | 0.0 | -0.5 | -2.9 | -261.1 |
| chr6B | 176131494 | TraesCS6B02G167300 | 11.9 | 9.0 | 0.0 | 2.4 | -2.9 | -262.5 |
| chr3B | 770556671 | TraesCS3B02G528300 | 9.9 | 7.0 | 0.0 | 0.2 | -2.9 | -262.8 |

| | | | | | | | | |
|---|---|---|---|---|---|---|---|---|
| chr3D | 589607809 | TraesCS3D02G500300 | 8.2 | 5.3 | 0.0 | -1.4 | -2.9 | -264.2 |
| chr5B | 488190034 | TraesCS5B02G304200 | 10.6 | 7.6 | 0.0 | 0.9 | -3.0 | -266.2 |
| chrUn | 382192747 | TraesCSU02G551500LC | 11.4 | 8.4 | 0.0 | 1.8 | -3.0 | -268.1 |
| chrUn | 87231917 | TraesCSU02G098500 | 8.6 | 5.6 | 0.0 | -1.1 | -3.0 | -268.2 |
| chr1D | 477502630 | TraesCS1D02G422100 | 10.0 | 7.0 | 0.0 | 0.2 | -3.0 | -270.4 |
| chr3D | 42395518 | TraesCS3D02G083600 | 13.0 | 9.9 | 0.0 | 3.6 | -3.0 | -273.0 |
| chr7B | 394912577 | TraesCS7B02G215400 | 7.9 | 4.8 | 0.0 | -1.7 | -3.1 | -278.1 |
| chr5B | 506656246 | TraesCS5B02G321100 | 11.5 | 8.4 | 0.0 | 2.0 | -3.1 | -278.6 |
| chr7A | 515199226 | TraesCS7A02G352000 | 12.3 | 9.2 | 0.0 | 2.8 | -3.1 | -278.8 |
| chr2B | 15880942 | TraesCS2B02G033000 | 10.3 | 7.1 | 0.0 | 0.7 | -3.1 | -280.4 |
| chr7B | 113444856 | TraesCS7B02G098900 | 7.9 | 4.8 | 0.0 | -1.7 | -3.1 | -281.5 |
| chr5B | 710325045 | TraesCS5B02G565400 | 9.7 | 6.5 | 0.0 | -0.1 | -3.2 | -283.6 |
| chr5A | 55248978 | TraesCS5A02G087100LC | 12.8 | 9.6 | 0.0 | 3.4 | -3.2 | -284.5 |
| chrUn | 387128454 | TraesCSU02G252000 | 7.8 | 4.6 | 0.0 | -1.8 | -3.2 | -284.5 |
| chr5A | 513639716 | TraesCS5A02G435400LC | 8.5 | 5.3 | 0.0 | -1.2 | -3.2 | -286.8 |
| chr4B | 24746070 | TraesCS4B02G033500 | 11.2 | 8.0 | 0.0 | 1.6 | -3.2 | -287.9 |
| chr4D | 14857918 | TraesCS4D02G031900 | 7.6 | 4.4 | 0.0 | -1.9 | -3.2 | -289.2 |
| chr2D | 9642976 | TraesCS2D02G020300 | 8.3 | 5.1 | 0.0 | -1.3 | -3.2 | -289.6 |
| chr1B | 646536082 | TraesCS1B02G711900LC | 8.6 | 5.4 | 0.0 | -1.1 | -3.3 | -294.1 |
| chrUn | 390206597 | TraesCSU02G253400 | 9.2 | 5.9 | 0.0 | -0.6 | -3.3 | -297.1 |
| chr1B | 677131140 | TraesCS1B02G467300 | 9.1 | 5.8 | 0.0 | -0.7 | -3.3 | -297.6 |
| chrUn | 184212574 | TraesCSU02G150700 | 11.9 | 8.5 | 0.0 | 2.3 | -3.4 | -302.5 |
| chr7A | 507013183 | TraesCS7A02G345000 | 8.3 | 4.9 | 0.0 | -1.4 | -3.4 | -303.9 |
| chr3A | 747991284 | TraesCS3A02G737400LC | 7.4 | 4.0 | 0.0 | -2.1 | -3.4 | -303.9 |
| chr1D | 442685785 | TraesCS1D02G359200 | 8.7 | 5.3 | 0.0 | -1.0 | -3.4 | -304.2 |
| chr3B | 828417030 | TraesCS3B02G609300 | 7.6 | 4.2 | 0.0 | -1.9 | -3.4 | -306.0 |
| chr5A | 699840782 | TraesCS5A02G544500 | 13.1 | 9.7 | 0.0 | 3.7 | -3.4 | -307.6 |
| chr4A | 677968336 | TraesCS4A02G404200 | 7.7 | 4.2 | 0.0 | -1.9 | -3.4 | -309.0 |
| chr7B | 505450448 | TraesCS7B02G276100 | 11.5 | 8.1 | 0.0 | 1.9 | -3.4 | -309.1 |
| chr6B | 30870897 | TraesCS6B02G067200LC | 8.9 | 5.5 | 0.0 | -0.9 | -3.4 | -309.7 |
| chrUn | 88203520 | TraesCSU02G099800 | 12.3 | 8.8 | 0.0 | 2.8 | -3.5 | -311.8 |
| chr1D | 477508260 | TraesCS1D02G422200 | 9.8 | 6.2 | 0.0 | -0.1 | -3.5 | -318.0 |
| chr3A | 405624910 | TraesCS3A02G219900 | 10.3 | 6.8 | 0.0 | 0.6 | -3.6 | -320.4 |
| chr1B | 669336052 | TraesCS1B02G452700 | 12.8 | 9.2 | 0.0 | 3.4 | -3.6 | -323.9 |
| chr7D | 479625402 | TraesCS7D02G370900 | 12.1 | 8.5 | 0.0 | 2.6 | -3.6 | -324.9 |
| chr7B | 682282185 | TraesCS7B02G414300 | 13.7 | 10.1 | 0.0 | 4.3 | -3.6 | -325.3 |
| chr2A | 696219175 | TraesCS2A02G447300 | 9.6 | 5.9 | 0.0 | -0.3 | -3.7 | -329.3 |
| chr2B | 93137930 | TraesCS2B02G124800 | 12.5 | 8.8 | 0.0 | 3.1 | -3.7 | -333.3 |
| chr5D | 78503927 | TraesCS5D02G110100LC | 7.4 | 3.7 | 0.0 | -2.1 | -3.7 | -335.6 |
| chr2B | 25413964 | TraesCS2B02G053000LC | 10.2 | 6.3 | 0.0 | 0.4 | -3.9 | -347.2 |
| chr4D | 486147812 | TraesCS4D02G326300 | 8.1 | 4.2 | 0.0 | -1.6 | -3.9 | -349.4 |
| chr7B | 122429103 | TraesCS7B02G105800 | 9.3 | 4.9 | 0.0 | -0.6 | -4.3 | -390.2 |
| chr1B | 16672853 | TraesCS1B02G033200LC | 13.3 | 8.9 | 0.0 | 3.8 | -4.4 | -392.6 |
| chr3D | 570311668 | TraesCS3D02G467400 | 9.3 | 4.9 | 0.0 | -0.6 | -4.4 | -393.7 |

| | | | | | | | | |
|---|---|---|---|---|---|---|---|---|
| chr5A | 589583883 | TraesCS5A02G394500 | 8.5 | -1.6 | 0.0 | 0.2 | -10.1 | -404.6 |
| chr2B | 408300285 | TraesCS2B02G461000LC | 13.1 | 8.5 | 0.0 | 3.6 | -4.6 | -418.3 |
| chr2B | 11391067 | TraesCS2B02G024000LC | 7.5 | 2.8 | 0.0 | -2.1 | -4.8 | -428.6 |
| chr2D | 579128018 | TraesCS2D02G589700LC | 7.7 | 2.8 | 0.0 | -2.0 | -4.9 | -440.1 |
| chr4B | 649353802 | TraesCS4B02G358600 | 7.8 | 2.8 | 0.0 | -1.9 | -5.0 | -449.1 |
| chr5A | 700372271 | TraesCS5A02G545200 | 13.5 | 8.3 | 0.0 | 4.0 | -5.2 | -469.1 |
| chrUn | 67081637 | TraesCSU02G120300LC | 10.5 | 5.2 | 0.0 | 0.6 | -5.3 | -475.2 |
| chr5A | 496334117 | TraesCS5A02G289800 | 8.2 | 2.8 | 0.0 | -1.6 | -5.4 | -490.3 |
| chrUn | 328173592 | TraesCSU02G222000 | 12.7 | 7.2 | 0.0 | 3.1 | -5.5 | -491.9 |
| chr7B | 617571591 | TraesCS7B02G357100 | 7.5 | 2.0 | 0.0 | -2.1 | -5.5 | -497.5 |
| chr5A | 699562234 | TraesCS5A02G724100LC | 13.2 | 7.7 | 0.0 | 3.7 | -5.6 | -500.0 |
| chr4B | 1080301 | TraesCS4B02G001900 | 8.4 | 2.8 | 0.0 | -1.5 | -5.6 | -503.3 |
| chr5A | 699630209 | TraesCS5A02G543900 | 12.1 | 6.4 | 0.0 | 2.4 | -5.6 | -506.0 |
| chr2D | 109129771 | TraesCS2D02G164900 | 9.0 | 3.3 | 0.0 | -1.0 | -5.7 | -511.3 |
| chr2B | 121376773 | TraesCS2B02G153000 | 10.0 | 4.2 | 0.0 | 0.1 | -5.8 | -519.3 |
| chr3B | 39344160 | TraesCS3B02G077100LC | 9.7 | 3.7 | 0.0 | -0.2 | -6.0 | -542.0 |
| chr3A | 9073442 | TraesCS3A02G011000 | 9.9 | 3.7 | 0.0 | 0.0 | -6.2 | -559.0 |
| chr7A | 12894244 | TraesCS7A02G035200LC | 8.2 | 2.0 | 0.0 | -1.6 | -6.3 | -563.5 |
| chr2B | 793971622 | TraesCS2B02G617400 | 8.3 | 2.0 | 0.0 | -1.7 | -6.3 | -566.6 |
| chr1D | 7694093 | TraesCS1D02G017000 | 8.6 | 2.0 | 0.0 | -1.3 | -6.7 | -599.9 |
| chr1A | 9156307 | TraesCS1A02G017300 | 8.8 | 2.0 | 0.0 | -1.2 | -6.8 | -611.7 |
| chr2A | 291642669 | TraesCS2A02G291300LC | 9.6 | 2.8 | 0.0 | -0.3 | -6.8 | -615.6 |
| chr2D | 566979655 | TraesCS2D02G460200 | 12.6 | 5.5 | 0.0 | 3.0 | -7.1 | -634.6 |
| chr3A | 51408022 | TraesCS3A02G079500 | 9.2 | 2.0 | 0.0 | -0.8 | -7.2 | -646.5 |
| chr5B | 655432065 | TraesCS5B02G693900LC | 7.7 | 0.0 | 0.0 | -2.0 | -7.7 | -694.7 |
| chrUn | 349092786 | TraesCSU02G494800LC | 11.4 | 3.7 | 0.0 | 1.8 | -7.8 | -697.7 |
| chr7B | 685269722 | TraesCS7B02G417400 | 7.8 | 0.0 | 0.0 | -1.9 | -7.8 | -705.4 |
| chr1A | 57217338 | TraesCS1A02G103100LC | 11.9 | 4.0 | 0.0 | 2.2 | -7.9 | -712.1 |
| chr6A | 6746558 | TraesCS6A02G016500LC | 9.9 | 2.0 | 0.0 | -0.1 | -8.0 | -715.6 |
| chr3A | 12117137 | TraesCS3A02G020800 | 8.1 | 0.0 | 0.0 | -1.7 | -8.1 | -730.6 |
| chr4B | 11995638 | TraesCS4B02G016600 | 8.1 | 0.0 | 0.0 | -1.7 | -8.1 | -733.0 |
| chr5B | 658826221 | TraesCS5B02G488700 | 10.3 | 2.0 | 0.0 | 0.4 | -8.3 | -749.0 |
| chr2B | 789669919 | TraesCS2B02G904700LC | 12.4 | 4.0 | 0.0 | 2.7 | -8.4 | -754.2 |
| chr5B | 531510313 | TraesCS5B02G350700 | 10.4 | 2.0 | 0.0 | 0.4 | -8.4 | -755.0 |
| chr5D | 172880292 | TraesCS5D02G177100LC | 9.0 | 0.0 | 0.0 | -1.1 | -9.0 | -806.8 |
| chr7B | 579825342 | TraesCS7B02G326300 | 9.0 | 0.0 | 0.0 | -1.0 | -9.0 | -813.1 |
| chr2B | 14600183 | TraesCS2B02G030800LC | 9.1 | 0.0 | 0.0 | -1.0 | -9.1 | -816.8 |
| chr6A | 564276199 | TraesCS6A02G332900 | 9.5 | 0.0 | 0.0 | -0.5 | -9.5 | -854.0 |
| chr2B | 15636853 | TraesCS2B02G032400 | 9.6 | 0.0 | 0.0 | -0.3 | -9.6 | -861.2 |
| chr7B | 685269137 | TraesCS7B02G700900LC | 9.6 | 0.0 | 0.0 | -0.4 | -9.6 | -861.4 |
| chr5D | 172857077 | TraesCS5D02G176900LC | 9.9 | 0.0 | 0.0 | -0.1 | -9.9 | -891.1 |
| chr5A | 699519934 | TraesCS5A02G543600 | 10.3 | 0.0 | 0.0 | 0.3 | -10.3 | -924.0 |
| chr4A | 2984490 | TraesCS4A02G004300 | 10.4 | 0.0 | 0.0 | 0.4 | -10.4 | -933.8 |
| chr1B | 100541434 | TraesCS1B02G096400 | 13.7 | 3.3 | 0.0 | 4.2 | -10.4 | -939.8 |

| | | | | | | | | |
|---|---|---|---|---|---|---|---|---|
| chr7B | 683030505 | TraesCS7B02G415300 | 10.7 | 0.0 | 0.0 | 0.9 | -10.7 | -964.4 |
| chr2A | 84233348 | TraesCS2A02G139000 | 10.8 | 0.0 | 0.0 | 0.8 | -10.8 | -969.2 |
| chr2B | 13716298 | TraesCS2B02G029800 | 11.7 | 0.0 | 0.0 | 2.0 | -11.7 | -1053.1 |
| chr7B | 746530047 | TraesCS7B02G827300LC | 12.0 | 0.0 | 0.0 | 2.3 | -12.0 | -1080.3 |
| chr6B | 4532464 | TraesCS6B02G006600 | 12.1 | 0.0 | 0.0 | 2.4 | -12.1 | -1091.5 |

*Rfc: gene expression log ratio of Robin under inoculation with water (control) or *Fusarium graminearum* (*Fg*) inoculum.

Tfc: gene expression log ratio of Tenacious under inoculation with water (control) or *Fusarium graminearum* (*Fg*)

TRc: gene expression log ratio of Tenacious versus Robin under inoculation with *water (control).*

logCPM3: log of average read counts per million of Tenacious versus Robin under inoculation with *Fusarium graminearum* (*Fg*)

TRf: gene expression log ratio of Tenacious versus Robin under inoculation with *Fusarium graminearum* (*Fg*)

DrGs = (Tfc- Rfc) /(|TRc| +0.01111)

Table S6. RNA-seq data were downloaded from NCBI and sample information.

| **accession** | **wheat cv** | **phenotype** | **Innoculation** | **reference** |
|---|---|---|---|---|
| SRS6616128 | AAC Tenacious | FHB resistance | Fg | Nilsen et al, 2021 |
| SRS6616138 | AAC Tenacious | FHB resistance | water | Nilsen et al, 2021 |
| SRS6616122 | Roblin | FHB susceptible | Fg | Nilsen et al, 2021 |
| SRS6616118 | Roblin | FHB susceptible | water | Nilsen et al, 2021 |
| SRR8885513 | AC Barrie | FHB resistance | None | Xiang et al, 2019 |
| SRR8885514 | AC Barrie | FHB resistance | None | Xiang et al, 2020 |
| SRR3090577 | Fielder | FHB susceptible | Fg | Buhrow et al, 2020 |

Table S7. Demonstration of data integration

| Br | Parent B (AC Barrie) | Parent A (Reeder) | variant sites | Br line1 | Br line2 | Br line3 | Br line4 | Br line5 |
|---|---|---|---|---|---|---|---|---|
| | GG | CC | 1 | GG(B) | GG(B) | CC(A) | GG(B) | CC(A) |
| | AA | GG | 2 | AA(B) | AA(B) | AA(B) | GG(A) | GG(A) |
| | AA | CC | 3 | AA(B) | CC(A) | CC(A) | CC(A) | AA(B) |
| | AA | CC | 4 | AA(B) | AA(B) | AA(B) | CC(A) | AA(B) |

| cB | Parent B (AC Barrie) | Parent A (Cutler) | | cB line1 | cB line2 | cB line3 | cB line4 | cB line5 |
|---|---|---|---|---|---|---|---|---|
| | GG | CC | 1 | GG(B) | CC(A) | GG(B) | GG(B) | CC(A) |
| | TT | GG | 2 | TT(B) | GG(A) | GG(A) | TT(B) | GG(A) |
| | AA | TT | 3 | TT(A) | AA(B) | TT(A) | TT(A) | AA(B) |
| | AA | CC | 4 | AA(B) | CC(A) | AA(B) | CC(A) | CC(A) |

| BrcB | | Br line1 | Br line2 | Br line3 | Br line4 | Br line5 | cB line1 | cB line2 | cB line3 | cB line4 | cB line5 |
|---|---|---|---|---|---|---|---|---|---|---|---|
| | 1 | GG(B) | GG(B) | CC(A) | GG(B) | CC(A) | GG(B) | CC(A) | GG(B) | GG(B) | CC(A) |
| | 3 | AA(B) | CC(A) | CC(A) | CC(A) | AA(B) | TT(A) | AA(B) | TT(A) | TT(A) | AA(B) |
| | 4 | AA(B) | AA(B) | AA(B) | CC(A) | AA(B) | AA(B) | CC(A) | AA(B) | CC(A) | CC(A) |

| BrcBa | | Br line1 | Br line2 | Br line3 | Br line4 | Br line5 | cB line1 | cB line2 | cB line3 | cB line4 | cB line5 |
|---|---|---|---|---|---|---|---|---|---|---|---|
| | 1 | GG(B) | GG(B) | CC(A) | GG(B) | CC(A) | GG(B) | CC(A) | GG(B) | GG(B) | CC(A) |
| | 2 | (A) | (A) | (A) | GG(A) | GG(A) | (A) | GG(A) | GG(A) | (A) | GG(A) |
| | 3 | AA(B) | CC(A) | CC(A) | CC(A) | AA(B) | TT(A) | AA(B) | TT(A) | TT(A) | AA(B) |
| | 4 | AA(B) | AA(B) | AA(B) | CC(A) | AA(B) | AA(B) | CC(A) | AA(B) | CC(A) | CC(A) |

| BrcBb | | Br line1 | Br line2 | Br line3 | Br line4 | Br line5 | cB line1 | cB line2 | cB line3 | cB line4 | cB line5 |
|---|---|---|---|---|---|---|---|---|---|---|---|
| | 1 | GG(B) | GG(B) | CC(A) | GG(B) | CC(A) | GG(B) | CC(A) | GG(B) | GG(B) | CC(A) |
| | 2 | (B) | (B) | (B) | GG(A) | GG(A) | (B) | GG(A) | GG(A) | (B) | GG(A) |
| | 3 | AA(B) | CC(A) | CC(A) | CC(A) | AA(B) | TT(A) | AA(B) | TT(A) | TT(A) | AA(B) |
| | 4 | AA(B) | AA(B) | AA(B) | CC(A) | AA(B) | AA(B) | CC(A) | AA(B) | CC(A) | CC(A) |

*GG(B): genotype (B, A, binary code for dpGLAS). B: AC Barrie, A: Reeder or Cutler

the integrated sets have the same number of sample number, but different number and different types of genotype variants.
